# Specialized layer 5 cortical terminals target functional thalamic spines

**DOI:** 10.64898/2026.07.05.735839

**Authors:** Hajnalka Bokor, Nóra Hádinger, Boglárka Tóth, Andor Domonkos, Eriko Kuramoto, Gordon Shepherd, Naoki Yamawaki, Judit Makara, László Acsády

## Abstract

Dendritic spines are ubiquitous morpho-functional elements of excitatory synaptic transmission and plasticity. Whether axo-spinous communication exists in the thalamus is presently unclear. Here we find that layer 5 (L5) corticothalamic terminals arising from frontal cortices selectively target thalamic spines with classical head and neck dimensions. The synapses of the L5 axo-spinous contacts were enriched in GluR1 subunit of AMPA receptor and their size correlated with their spine head volume. Synaptic stimulation of thalamic spines resulted in compartmentalized Ca^2+^ responses similar to that of hippocampal spines. Optogenetic activation of the frontal L5-thalamus pathway had strong impact on thalamic spiking even at high stimulation frequencies and could selectively recruit a subnetwork of thalamic cells in a behavioral state dependent manner. Optogenetic interference with the small L5-thalamic pathway perturbed motor learning. Our study reveals a hitherto unrecognized, effective top-down communication channel between the frontal cortex and thalamus that involves highly variable axo-spinous contacts.

---

Dendritic spines are major components of excitatory synaptic transmission^1–4^. The vast majority of glutamatergic synaptic communication in the cortex takes place via axo-spinous connections^5,6^. Cortical spines are highly variable in size and shape^4,7^. They typically display compartmentalized Ca^2+^ responses upon synaptic activation, which is a major determinant of input-specific, activity-dependent alterations in synaptic strength^8^. This mechanism is thought to be fundamental for the encoding of novel information in the nervous system manifested by the flexible and selective recruitment of distinct neuronal populations^9,10^.

Spines were described on neurons in the thalamus as well by Ramon y Cajal over 100 years ago^11^. Later, thalamic spines were also used to classify thalamic neurons^12^. However, the ultrastructure, the source of afferent inputs and the functional properties of thalamic spines remained obscure. It is also presently unclear if postsynaptic Ca^2+^ entry follows synaptic activation in thalamic spines^13^. This leaves the questions open whether they operate equivalently to cortical spines and whether they may serve as a specialized target for certain thalamic input pathway(s). Since thalamus is involved in all cortical functions and dysfunctions via the complex thalamo-corticothalamic excitatory communication system^14^, understanding the basics of synaptic communication via spines is key to resolve thalamic information transfer.

Potential excitatory afferents to thalamic spines are numerous since the input space of thalamus involves almost the entire neuraxis including subcortex and cortex^15^. Subcortical glutamatergic inputs to the thalamus mostly form giant terminals 3-8 µm in diameter^15–17^ establishing multisynaptic contacts with complex, branching dendritic appendages of thalamic neurons^17,18^. These appendages are fundamentally different from cortical spines that are known to receive mostly single synapses^4,5,19^. Cortical glutamatergic inputs to the thalamus have two distinct components, one originating in layer 5 (L5) and the other in layer 6 (L6)^20–24^. The L5 corticothalamic pathways, studied so far, have been shown to establish large multisynaptic terminals identical to the subcortical large terminals both in morphology and physiology: they target complex dendritic appendages, not classical spines^23,25–28^. The L6 corticothalamic terminals are small and innervate mainly dendritic shafts^29–31^, leaving the major source of the inputs to thalamic spines unknown.

The properties of the L5 and L6 corticothalamic inputs listed above have been mainly based on the investigation of sensory cortices. The precise morphology and physiology of the corticothalamic terminals arising from the frontal cortices have received much less attention^32–36^ even though frontal cortex has an active role in recruiting thalamic neurons during complex cognitive functions that require flexible allocation of thalamic neuronal populations^37–39^. Recently, frontal cortical L5 afferents have been shown to form powerful connections with the thalamic reticular nucleus (TRN), the major inhibitory interface of the thalamus^40^. However, similar studies aiming to investigate the morphological and physiological features of the frontal L5 pathways to the thalamus are lacking.

To fill this gap in our knowledge here we studied L5 corticothalamic terminals arising from frontal cortical regions. We found that they are significantly different both from sensory cortical L5 and frontal L6 terminals. Unlike sensory L5, frontal L5 formed only small terminals in the thalamus which preferentially targeted functional spines via variable, complex but mostly single synapses. The axo-spinous L5-thalamic synaptic connections were able to selectively recruit subsets of thalamic neurons and were involved in motor learning.

## Results

### Axo-spinous connections between cortical L5 and the thalamus

For a comparison of the sensory and frontal L5-thalamic pathways we selected a prototypical sensory cortical region, the S1 somatosensory cortex, and compared it to the secondary M2 motor cortex, a frontal isocortical region known to be involved in higher order motor and posture control^41,42^. We injected AAV-dio-ChR2-EYFP into S1 and M2 of Rbp4-Cre mice to selectively label L5 corticothalamic axons^40^ and first examined the morphology of the terminal boutons in their main thalamic target nuclei, the posterior nucleus (Po) and the ventromedial nucleus (VM), respectively (Fig. 1a, Extended Data Fig. 1). Using high power confocal microscopy, initial visual inspection indicated a sharp contrast between the two pathways. Axons from the S1/L5 to Po (S1/L5-Po) formed many large sized terminals (>∼3 µm) as described before^26,28,43,44^(Fig. 1b, top). However, such large axon terminals were not observed in the M2/L5 to VM (M2/L5-VM) pathway; instead, the boutons were universally small (∼1 µm) (Fig. 1b, bottom). For a quantitative comparison we measured the largest cross-sectional area (CSA) of boutons using Z-stacks of high-power confocal images. The CSAs of S1/L5-Po terminals were skewed to higher values due to the presence of large-sized terminals, whereas the CSAs of VM terminals never exceeded 1 µm^2^ and were significantly smaller than S1/L5-Po boutons (Fig. 1c). These data show that L5 corticothalamic terminals display a pronounced region-specific difference in size, as suggested before ^35^, and that VM is contacted only by small L5 terminals from M2.

**Figure 1.**
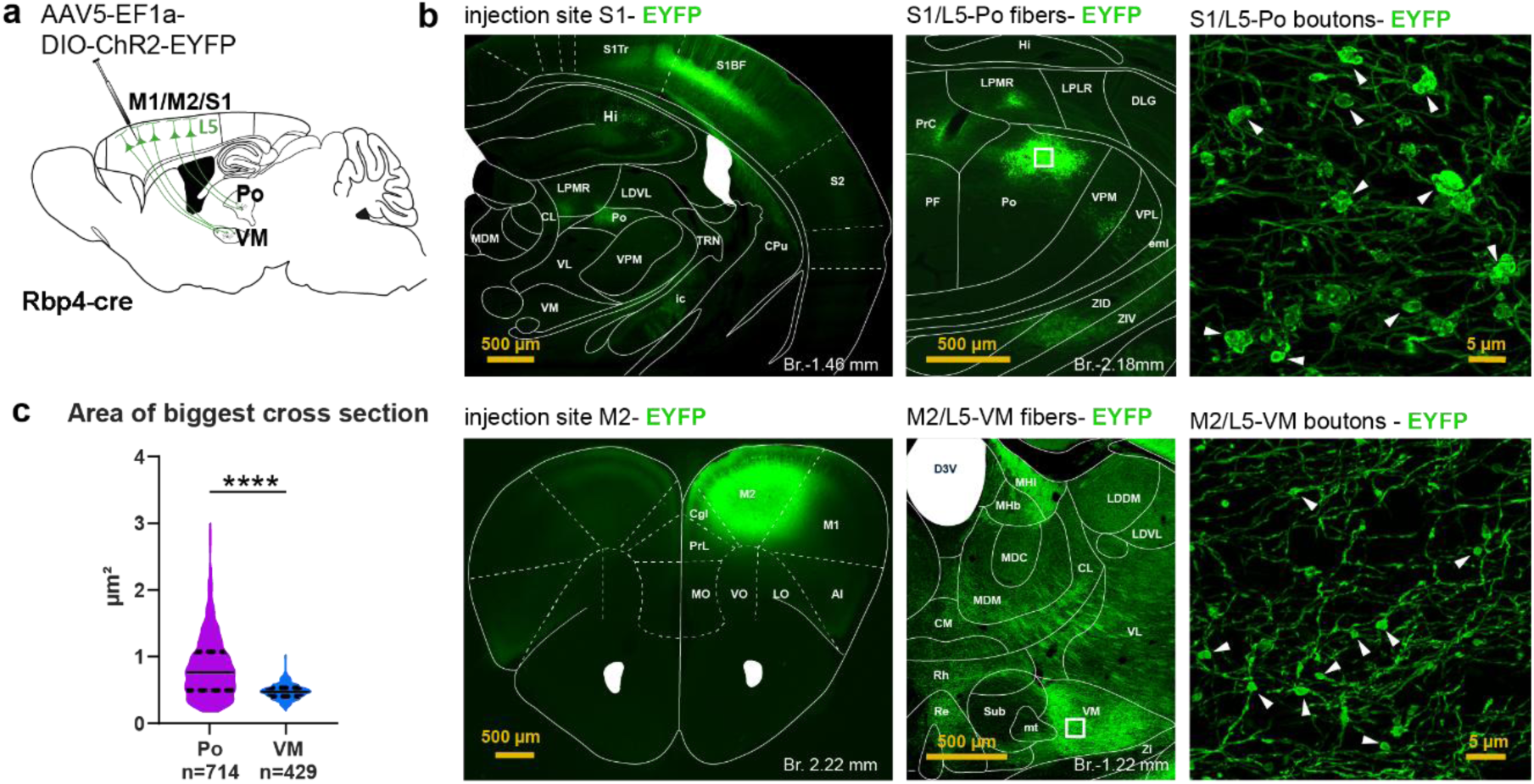
Absence of large terminals in the M2/L5-thalamic pathway. a) Experimental design b) Extent of S1 (top) and M2 (bottom) injection sites (left) and termination fields in the thalamus (middle) and high power images of L5 axon terminals (right, arrowheads) from the area marked with white rectangles. c) Size distribution of M2/L5-VM and S1/L5-Po axon terminals (Mann-Whitney test, p <0.0001)

The large S1/L5-Po terminals are known to form multisynaptic contacts on the dendritic appendages of thick proximal dendrites^26,43–45^. To examine the synaptic organization of the small M2/L5 terminals, we performed 3D electron microscopic reconstructions of labeled L5 terminals in the VM. Since L6 terminals are also small we compared the ultrastructure of the M2/L5 terminals to that of the M2/L6 terminals in VM. L5 boutons from the M2 cortex were labeled in Rbp4-Cre mice (n=61 L5 boutons in n=2 mice, Fig. 2a; the injection sites in both cases involved also regions of M1 next to M2 but did not spread to S1). For the study of L6 boutons we used two approaches in two different mouse strains, since not all AAV constructs label L6 well^46^. First, we iontophorized Phaseolus vulgaris leucoagglutinin (PHAL) targeting the L6 in a Thy1-ChR2-EYFP mice, where L5 cells express EYFP (n=24 boutons in n=1 mouse, Extended Data Fig. 2a, b). In this animal PHAL- and Thy-1 EYFP-positive terminals did not overlap in the VM (Extended Data Fig. 2b) demonstrating selective labeling of L6 boutons by PHAL. Second, in another animal we injected AAV-dio-ChR2-EYFP into the M2 of an Ntsr1-Cre mouse (Fig. 2b, Extended Data Fig. 2c, n=12 boutons in n=1 mouse). Since the properties of the L6 boutons labeled by the two methods did not differ significantly (Extended Data Fig. 2d-i), we pooled the two datasets.

**Figure 2.**
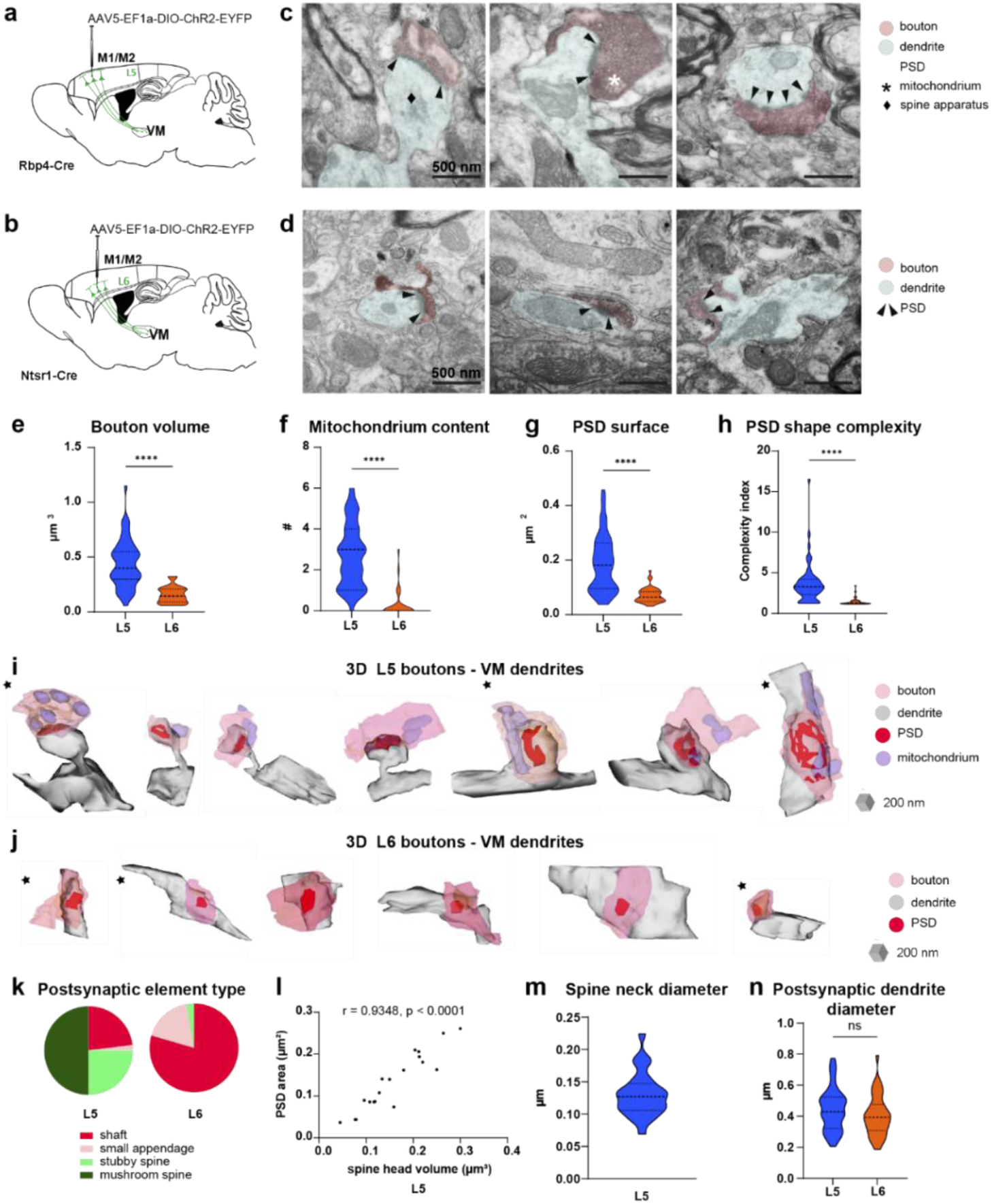
Small M2/L5 boutons target dendritic spines in the thalamus unlike small M2/L6 terminals. a-b) Experimental designs. c) Electron micrographs of L5 terminals (pink shading) establishing synapses on a mushroom spine (left), a stubby spine (middle) and a dendritic shaft (right) in VM (blue shading). For 3D reconstructions see panel i. d) Electron micrographs of L6 terminals (pink shading) establishing synapses on dendritic shafts (blue shading, left and middle) and on a small dendritic appendage (right). Black arrowheads: PSD. For 3D reconstructions see panel j. e) Bouton volume (L5: 0.44±0.03 µm^3^ n=52 in n=2 mice; L6: 0.16±0.02 µm^3^ n=24 in n=2 mice; Unpaired t test p<0.0001). f) Number of mitochondria (L5: 2.71±0.22 n=49 in n=2 mice; L6: 0.3±0.14 n=27 in n=2 mice; Mann Whitney test: p<0.0001). g) PSD surface (L5: 0.19±0.01 µm^2^ n=42 in n=2 mice; L6: 0.07±0.01 µm^2^ n=28 in n=2 mice; Mann Whitney test: p<0.0001). h) PSD shape complexity (L5: 3.73±0.37 n=49 in n=2 mice; L6: 1.43±0.09 n=31 in n=2 mice; Mann Whitney test: p<0.0001). i) Examples for 3D reconstructed L5 boutons (pink, mitochondria in purple) in the VM and their postsynaptic targets (grey, PSD in red): mushroom spines (left 1-4), stubby spines (middle 5-6), or a dendritic shaft (right 7). Black asterisks label structures shown in (c). j) Examples for 3D reconstructed L6 boutons (pink, mitochondria in purple) in the VM and their postsynaptic targets (grey, PSD in red): dendritic shafts (left 1-5) or a small appendage (right 6). Black asterisks label structures shown in (d). k) Postsynaptic element type (L5: shaft 23.08%, small appendage: 1.92%, stubby spine: 25%, mushroom spine: 50%, n=52; L6: shaft 79.5%, small appendage 17.9%, stubby spine 2.6%, mushroom spine 0%, n=39) . l) Correlation of M2/L5-VM PSD area and spine head volume (L5: n=20 pairs in n=2 mice, Pearson correlation: r=0.9348, p<0.0001). m) Spine neck diameter (L5: 0.13±0.01 µm, n=26 in n=2 mice) n) Postsynaptic dendrite diameter (L5: 0.51±0.06 µm, n=45 in n=2 mice; L6: 0.40±0.02 µm, n=37 in n=2 mice; Mann Whitney test: ns). A single outlier (L5: 3.184 µm) is not shown on the graph for clarity but is included in the analysis. Violin plots: horizontal endpoints label minimum and maximum values, scattered lines label first to third quartiles, dashed line labels median value. PSD: postsynaptic density.

Both M2/L5 and M2/L6 terminals established classical asymmetrical synapses in the VM (Fig. 2c,d). At the electron microscopic level M2/L5 boutons were small (0.44±0.03 µm^3^, n=52) confirming the confocal data but had a significantly larger volume than M2/L6 terminals (mean 0.16±0.02 µm^3^, n=24) (Fig. 2e). Almost all M2/L5 boutons in VM (48/49, 98%) contained one or multiple mitochondria, whereas mitochondria were missing in most M2/L6 boutons (22/27, 81%) (Fig. 2f). Multiple puncta adhaerentia, a characteristic of large L5 terminals arising from sensory cortices, were not observed at M2/L5 terminals. The area of postsynaptic densities (PSD) in case of M2/L5 terminals spanned a wide range (0.037 to 0.485 µm^2^ mean: 0.19 ±0.01 µm^2^, n=42) and was significantly larger than that of the M2/L6 terminals (0.07 ±0.01 µm^2^, n=28) (Fig. 2g). The shape of M2/L5 PSDs in VM was more complex than that of M2/L6 PSDs (Fig. 2h) and included perforated, branching or segmented synapses, unlike M2/L6 terminals that were always macular. With one exception, all M2/L5 terminals (n=49) targeted a single postsynaptic structure. Besides the ultrastructural features of the terminals, M2/L5 and M2/L6 terminals also showed a striking difference in postsynaptic target selectivity (Fig. 2i-k, Extended Data Fig. 2j). Most small M2/L5 boutons (39/52, 75%) targeted dendritic protrusions with classical spine morphology (Fig. 2c,i,k), in which spine apparatus could be frequently identified (73.53%, 25/34 in n=2 mice) (Fig. 2c). M2/L5 terminals that did not innervate spines targeted dendritic shafts (12/52, 23.08%) or in a single case, a small appendage (1/52, 1.92%) (Fig. 2i). The shape of the targeted spines was highly variable and ranged from mushroom spines with large head and thin neck (n=26/39) to stubby spines with little or no discernable neck (n=13/39) (Fig. 2i,k). In contrast, M2/L6 boutons established synapses primarily onto dendritic shafts (31/39, 79.5%) or small dendritic protrusions not exceeding 0.3 µm CSA (7/39 17.9%) which displayed no classical spine morphology (Fig. 2d,j,k, Extended Data Fig. 2i,j). A single M2/L6 bouton targeted a stubby spine (1/39 2.6%). The PSD area of M2/L5-spine synapses significantly correlated with the spine head volume (Fig. 2l) similarly to that of hippocampal spines ^47^. The spine neck diameter (0.13±0.01 µm, n=26, Fig. 2m) was also comparable to those of the hippocampal spines ^47^. We measured the diameter of the parent dendrites that gave rise to spines postsynaptic to small M2/L5 terminals as well as the diameter of postsynaptic shafts targeted by M2/L5 and M2/L6 terminals. Both pathways contacted primarily small to medium diameter dendrites (< 0.8 µm) that were not significantly different between M2/L5 and M2/L6 (L5: 0.51±0.06 µm n=45; L6: 0.40±0.02 µm, n=37, Fig. 2n). Since dendritic diameter decreases with distance from the soma in thalamic neurons^44,48^ these results indicate that the two terminal types target similar dendritic domains. Indeed, in the vicinity of labeled L5 axo-spinous contacts we often found small, putative L6 terminals without mitochondria establishing synapses on dendritic shafts.

In summary, these data show that small M2/L5 terminals display a unique pre- and postsynaptic organization, preferentially innervate dendritic spines and differ from both the small M2/L6 corticothalamic boutons and the known organization of large S1/L5 terminals^26,43–45^.

### Thalamic spines display compartmentalized Ca^2+^ responses

The ultrastructural evidence demonstrated that a large proportion of small-sized M2/L5 terminals in the VM target dendritic spines that display similar basic morphological features to cortical spines. However, it is presently unclear whether they can function as “real” spines; i.e., able to display compartmentalized Ca^2+^ response upon synaptic activation. We therefore investigated the functional properties of thalamic spines using patch-clamp recordings combined with two-photon (2P) Ca^2+^ imaging and glutamate uncaging (2PGU) in *in vitro* acute slice experiments (Fig. 3a).

**Figure 3.**
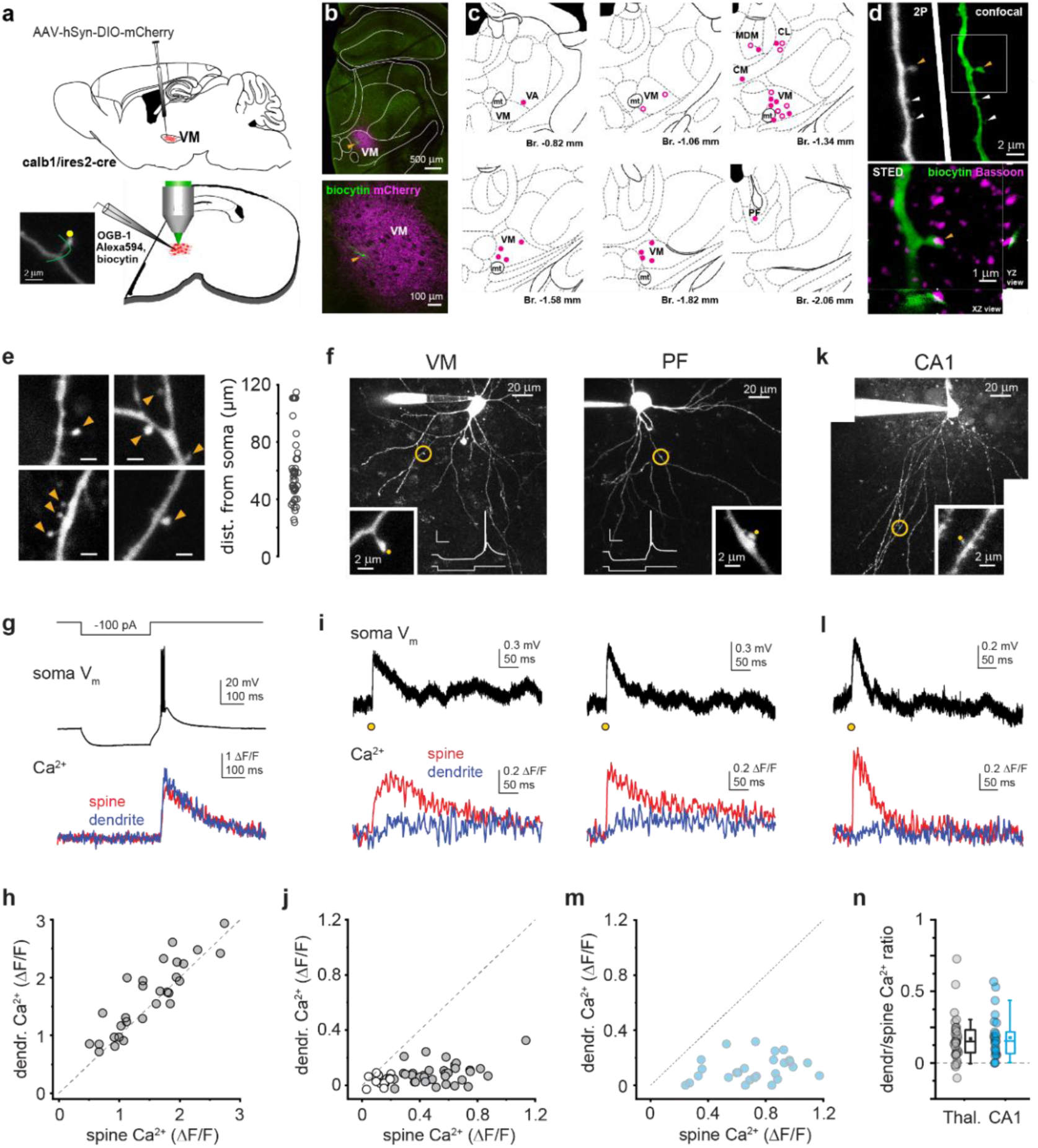
Thalamic spines compartmentalize synaptically evoked Ca^2+^ signals. a) Experimental design. Green line in the inset shows the path of the line scan. b) Representative AAV injection site in VM (upper panel, magenta) and a higher power confocal image of the imaged and biocytin labeled cell in VM (lower panel, green cell and orange arrowhead). c) Anatomical position of the recorded cells within the thalamus. Filled circles: 2PGU-tested cells in final analysis, open circles: additional cells with large spines. d) Upper panel: A representative spiny dendritic segment of a recorded cell in VM imaged with 2P (left) and *post hoc* with confocal microscope (right). Arrowheads point to spines. Orange arrowhead: A spine shown also on the lower panel in a STED image overlaid on Bassoon immunolabeling (confocal) of the presynaptic active zone on the spine head. e) Example 2P z-stack maximum projection images of spines in thalamic neurons. Right, Distance of spines from the soma. f) Two representative neurons investigated functionally in the VM (left) and PF (right). Yellow circles, the location of the stimulated spine shown in the insets at high power. Yellow dots, the location of 2PGU. g) Representative rebound spike in a patched VM thalamic neuron (cell and spine shown in panel f, left). Top, current injection protocol; middle, somatic voltage trace; bottom, Ca^2+^ recording in the spine and dendrite. h) Ca^2+^ signals evoked by rebound spikes in the spine head and dendritic shaft (n=29 spines). Dendrite/spine Ca^2+^ signal amplitude ratio for rebound spikes was larger than 1 (mean ± SEM: 1.15 ± 0.05, Wilcoxon test comparison to 1: p=0.007). i) Responses of the example spines in the neurons shown in panel f, to stimulation by 2PGU. Top, average somatic voltage response; bottom, average spine and dendrite Ca^2+^ response to 2PGU stimulation (yellow dots in panel f insets). j) Uncaging-evoked Ca^2+^ signals in the thalamic spine heads and dendritic shafts (n=49 stimulus series in n=27 spines). Grey filled dots show stimuli evoking >0.2 ΔF/F spine Ca^2+^ signal. k) An example spine stimulated by 2PGU in a basal dendrite of a CA1PC (2P z-stack image, with the location of the stimulated spine (inset, single scan) indicated by yellow circle. l) Average somatic voltage response (top) and average spine and dendrite Ca^2+^ response (bottom) to 2PGU stimulation (yellow dot in panel k inset) in the same CA1PC. m) Uncaging-evoked Ca^2+^ signals in the spine heads and dendritic shafts of CA1PCs (blue). n) Dendrite/spine Ca^2+^ signal amplitude ratio for spines with >0.2 ΔF/F Ca^2+^ signal (CA1PC spines: n=27, stimulation series in n=21 spines in 5 cells; thalamic neurons: n=36, stimulus series in n=24 spines in 16 cells; Mann-Whitney test: p=0.906). Box plots show mean (square), median (line), 25-75% (box) and 10-90% of the data (whiskers).

To identify the area of interest in the slice, we injected a Cre-dependent cytosolic mCherry-expressing AAV construct (n=35 animals, Fig. 3a) to the VM (Extended Data Fig. 3a) and the neighboring regions that were rich in small L5 terminals, of adult calb1/ires2cre mice. These nuclei contain many Calb1+ neurons^48^. After slice preparation, the region with mCherry-positive cells was identified by 2P imaging (Fig. 3b) and neurons in the area were selected for somatic whole-cell patch-clamp recording. *Post hoc* histology verified that all cells included in the analysis below were located in Calb1+ thalamic nuclei that are known to receive frontal cortical L5 innervation (Fig. 3b, c). The patched cells were loaded with the Ca^2+^ sensitive dye Oregon Green BAPTA-1 (OGB-1, 100 μM) and the structural fluorophore Alexa Fluor 594 (50 μM), allowing unambiguous visualization of their dendritic processes (Fig. 3d-f). Similar to the electron microscopic data, spine size and shape were highly variable in the *in vitro* material (Fig. 3d,e, Extended Data Fig. 3b). In addition to very small spine-like protrusions that were difficult to resolve with 2P imaging, in a subset (64%) of the recorded thalamic cells we could identify dendritic spines with a relatively large, well-defined head that was clearly separated from the dendritic shaft, and in many cases the spine neck could also be visualized (Fig. 3d-f). These large spines were sparse and typically located within 120 µm from the soma (Fig. 3e) and were used for further investigation. Some of the recorded spines were also identified *post hoc* using confocal and STED microscopy and were shown to contain synaptic markers of presynaptic density on the spine head (n = 6 cells, Fig. 3d, Extended Data Fig. 3b).

Recorded thalamic cells had resting V_m_ of -57.2 ± 0.5 mV (n=70) and R_in_ of 101 ± 11 MOhm (measured in n=18 cells) after break-in, and as expected, after the end of hyperpolarizing current injections (-50-200 pA for 300 ms) they all fired characteristic low-threshold rebound bursts (Fig. 3g). Ca^2+^ signals evoked in the recorded spines of thalamic cells during rebound spikes were uniformly large with slightly higher amplitudes in the dendritic shaft than in the spine head (n=29 spines in 21 cells, Fig. 3g-h), consistent with dendritic spikes propagating to the spines and inducing Ca^2+^ influx via voltage gated Ca^2+^ channels^49^.

To mimic synaptic inputs, we stimulated thalamic spines by 2P uncaging of MNI-glutamate (Fig. 3f,i, Extended Data Fig. 3c). Brief (0.2-0.5 ms) pulses of 720 nm uncaging laser light evoked EPSPs with fast rise (1.95 ± 0.24 ms 20-80% rise time, n=27 spines in 18 cells) as measured at the soma, with laser power adjusted to achieve reliable responses with moderately large amplitudes (<∼1.5 mV, mean ± SEM: 0.71 ± 0.05 mV; Extended Data Fig. 3c, d). EPSPs were strongly reduced by NBQX (10-20 μM applied in the bath, Extended Data Fig. 3e) indicating that they were mediated by AMPA receptors activated by glutamate, consistent with the presence of GluR subunits in VM^50^ (also see below). In the majority of spines, uncaging-evoked EPSPs were accompanied by sizable (>0.2 peak ΔF/F) Ca^2+^ transients in the spine head (Fig. 3i-j) that scaled with larger EPSPs (Extended Data Fig. 3d). In contrast, almost no response could be measured in the dendritic shaft (Fig. 3i-j), indicating strong compartmentalization of synaptically evoked Ca^2+^ signals in the spine head.

To compare this compartmentalization to that of spines with well-characterized function, we also measured Ca^2+^ transients in spines located on basal and oblique dendrites of hippocampal CA1 pyramidal cells (CA1PCs), a prototypical cortical cell type in which the morpho-functional properties of the spines are well known^51–53^ using the same brains (Fig. 3k-l). While the peak Ca^2+^ signal amplitudes corresponding to similar sized EPSPs were slightly but significantly smaller in thalamic spines (Extended Data Fig. 3f), the level of compartmentalization (quantified as the dendrite/spine Ca^2+^ signal amplitude ratio) by the spines responding with >0.2 ΔF/F Ca^2+^ signals was not significantly different in thalamic neurons and CA1PCs (Fig. 3j,m,n, thalamus: n=36 stimulation series in 24 spines in 16 cells; CA1PC: n=27 stimulation series in 21 spines in 5 cells; p=0.906, Mann-Whitney test).

Altogether, these results demonstrate that dendritic spines of thalamic neurons represent functionally isolated biochemical compartments for synaptic inputs similar to the spines of CA1 pyramidal cells.

### Cortical drive of thalamic activity via small L5 terminals in vivo

According to previous studies, reliable spike transmission in the thalamus is associated with inputs from large-sized multisynaptic, so called “driver” boutons^24,54^. In order to examine the effects of the small L5 corticothalamic terminals on the postsynaptic firing and to compare the influence of large vs. small L5 terminals on thalamocortical cells, we injected AAV5-EF1a-DIO-ChR2-EYFP into the S1 (n=5 animals) or M2 (n=18) cortices of Rbp4-Cre animals. We recorded the activity of postsynaptic Po (n=21) and VM (n=51) cells i*n vivo* following cortical laser stimulations. We used the juxtacellular recording and labeling approach^55^ in mice anesthetized with ketamine-xylazine, since this method allows unambiguous assignment of the recorded cells to a given thalamic nucleus as well as the visualization of the spatial relationship between the L5 afferents and the recorded cells (Fig. 4a,b, Extended Data Fig. 4a). Cortical activation evoked reliable, short latency (<20 ms) response in both the S1/L5-Po (n=8, Fig. 4c) and the M2/L5-VM (n=28, Fig. 4d) pathways in neurons that were found among the ChR2-EYFP labeled L5 fibers in the thalamus after the *post hoc* histological analysis (Fig. 4a-e). Neurons that were located outside of the labeled L5 axon arbor remained unresponsive to the cortical activation (n=4 identified cells in 4 animals, Extended Data Fig. 4a). Comparison of the response probabilities (See Methods) at the population level demonstrated that at 1 mW stimulation intensity and 1 Hz stimulation frequency VM cells could be activated at significantly higher probabilities than Po cells (Fig. 4f, Extended Data Fig. 4b). Using 4, 10 and 20 Hz stimulus trains, the firing probability decreased relative to the 1 Hz activation in both pathways at all frequencies tested (Fig. 4f, Extended Data Fig. 4b), but the response probabilities of the M2/L5-VM pathway remained significantly higher, especially at higher stimulation frequencies (Fig. 4f). Next, we compared the changes in response probabilities within the 10 Hz trains (10 stimuli) between the two pathways. At the start of the 10 Hz train both pathways displayed pronounced depression, however, during the later phases of the 10Hz trains the M2/L5-VM pathway showed a partial recovery whereas response probabilities of the S1/L5-Po pathway remained very low (Fig. 4g).

**Figure 4.**
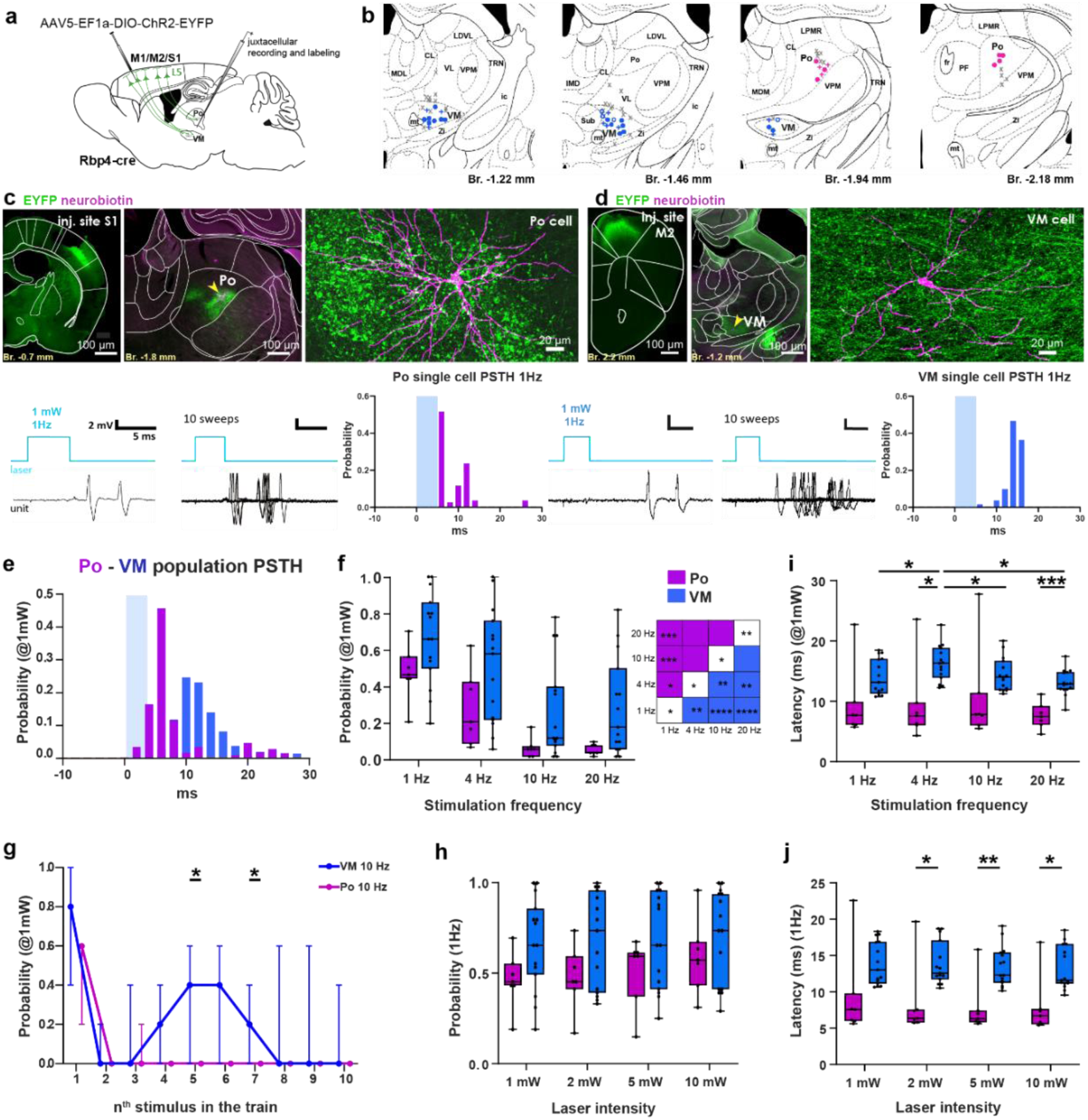
Optogenetic activation of thalamic cells by large S1/L5 and small M2/L5 terminals in vivo. a) Experimental design. b) Location of recorded neurons in VM (blue filled circles; soma and dendrites reconstructed, empty circles: only soma found) and Po (magenta circles; same as for blue). Colored +, responsive cells, not in the full analysis, Gray X, non-responsive cells. c) (top) Injection site in S1, low and high power images of S1/L5 large terminals in Po surrounding a representative juxtacellularly recorded and labeled Po cell. (bottom) Response of the same Po cell to the optogenetic stimulation of S1/L5. 1, 10 sweeps and PSTH. d) Same for a representative VM cell innervated by small M2/L5 terminals. e) Population PSTHs to cortical stimulation for Po (magenta) and VM (blue). f) Response probabilities of the two pathways for different stimulation frequencies. (2-way repeated measures (RM) ANOVA with Geisser-Greenhouse (GG) correction and followed by Tukey’s *post hoc* tests, interaction: F(3,60) = 0.1158, p=0.8479, (GG ε = 0.5348), frequency: F(3,60) = 43.80, p<0.0001, (GG ε = 0.2146), nucleus: F (1,20) = 5.050, p=0.0361. g) Median in train response probabilities of Po (magenta) and VM (blue) cells for 10 Hz stimulation (2-way RM ANOVA with GG correction, followed by Sidak’s multiple comparison tests, interaction: F(9,180) = 0.3941, p=0.6699 (GG å = 0.2146), stimulus in train: F(9,180) = 13.29, p<0.0001, (GG ε = 0.2146), nucleus: F (1,20) = 5.396, p=0.0308. h) Response probabilities for different stimulation intensities (2-way RM ANOVA with GG correction, as none of the ANOVA tests were significant, no *post hoc* test is shown, interaction: F(3,60) = 0.82, p=0.4605, (GG ε = 0.7723), intensity: F(3,60) = 2.35, p=0.0993, (GG ε = 0.7723), nucleus: F(1,20) = 2.80, p=0.1098. i-j) Response latencies for different stimulation frequencies I; 2-way RM ANOVA with GG correction and followed by Tukey’s *post hoc* tests, interaction: F(3,60) = 1.543, p=0.2265), (GG ε = 0.6586), frequency: F(3,60) = 4.265, p=0.0214, (GG ε = 0.6586), nucleus: F(1,20) = 10.84, p=0.0036, and intensities j; 2-way RM ANOVA with Geisser-Greenhouse correction and followed by Tukey’s *post hoc* tests, interaction: F(3,60) = 0.94, p=0.3927), (GG ε = 0.6023), intensity: F(3,60) = 4.32, p=0.0240, (GG ε = 0.6023), nucleus: F(1,20) = 11.31, p=0.0031,.\**P* < 0.05, \*\**P* < 0.01, \*\*\**P* < 0.001

Recruiting larger cortical input population by increasing laser power at 1 and 4 Hz (2, 5, 10 mW)^40^ did not result in significantly higher response probabilities in Po and VM (Fig. 4h, Extended data Fig. 4c). At 10 and 20 Hz stimulation, response probabilities of VM cells were significantly higher than those for Po at all laser intensities tested (Extended Data Fig. 4c). Response latencies were significantly longer in VM cells compared to Po for most stimulation frequencies and laser intensities tested (Fig. 4i,j, Extended Data Fig. 4d).

In a subset of cells response probabilities were tested from very low to high laser intensities (0.01 mW to 10 mW, VM, n=7, Po n=4, Extended Data Fig. 4e). The intensity/probability slopes for Po and VM were significantly different (simple linear regression: p= 0.0175).

These data together show that small M2/L5 terminals are able to drive postsynaptic action potentials in VM neurons with high (but variable) fidelity even at very low laser intensities. Furthermore, unlike the large S1/L5 terminals in Po, the M2/L5-VM pathway displayed high response probabilities even at higher stimulation frequencies.

### Small M2/L5 terminals display less short-term depression than large S1/L5 terminals

We found that in the intact animals the S1/L5 pathway (that has large terminals) displays pronounced short term depression (STD) (Fig. 4) in agreement with the literature^56^. On the other hand, small M2/L5 terminals evoked high-probability firing even at high presynaptic activity. To directly compare the short-term plasticity of the two types of L5 connections, we recorded thalamic postsynaptic EPSCs *in vitro* following photostimulation of L5 axons in the S1-Po and M2-VM pathways.

L5 afferents from S1 or M2 were virally labeled by injecting Cre-dependent AAV-DIO-ChR2 into Rbp4-Cre mice. After preparing the thalamic slice and establishing the whole-cell recording in a neuron, we first obtained a map of locations where photostimulation of the cortical axons evoked synaptic inputs, using laser-scanning photostimulation mapping (1 mW laser intensity). We then selected one of these sites that was located sufficiently far away from the soma and dendrites to ensure that the synaptic inputs recorded postsynaptically were not contaminated by artifactual “over-bouton” stimulation effects^57^. At this site, we applied a small-spot (∼50 µm) repetitive laser stimulation protocol with the laser intensity set to a minimal level that was sufficient to reliably evoke a response (Fig. 5a-d, Extended Data Fig. 5). We applied a 10-pulse stimulation at four different frequencies: 1, 5, 10, and 20 Hz (Fig. 5e). The mean response latency for the first evoked response in the power series experiments was 3.4 ms (n=7, SD: 0.54 ms, range, 2.7-4.1 ms) for Po and 4.4 ms (n=8, SD: 1.06 ms, range 3.2-6-2 ms) for VM cells (Fig. 5f). Both the S1/L5–Po and the M2/L5–VM pathways showed STD at all frequencies. However, the degree of STD was significantly less in the M2/L5-VM pathway compared to the S1/L5-Po pathway at 5, 10, and 20 Hz (Fig. 5g). These data can underlie the observed differences in the short term plasticity of Po and VM cells upon high frequency activation of L5 terminals *in vivo*.

**Figure 5.**
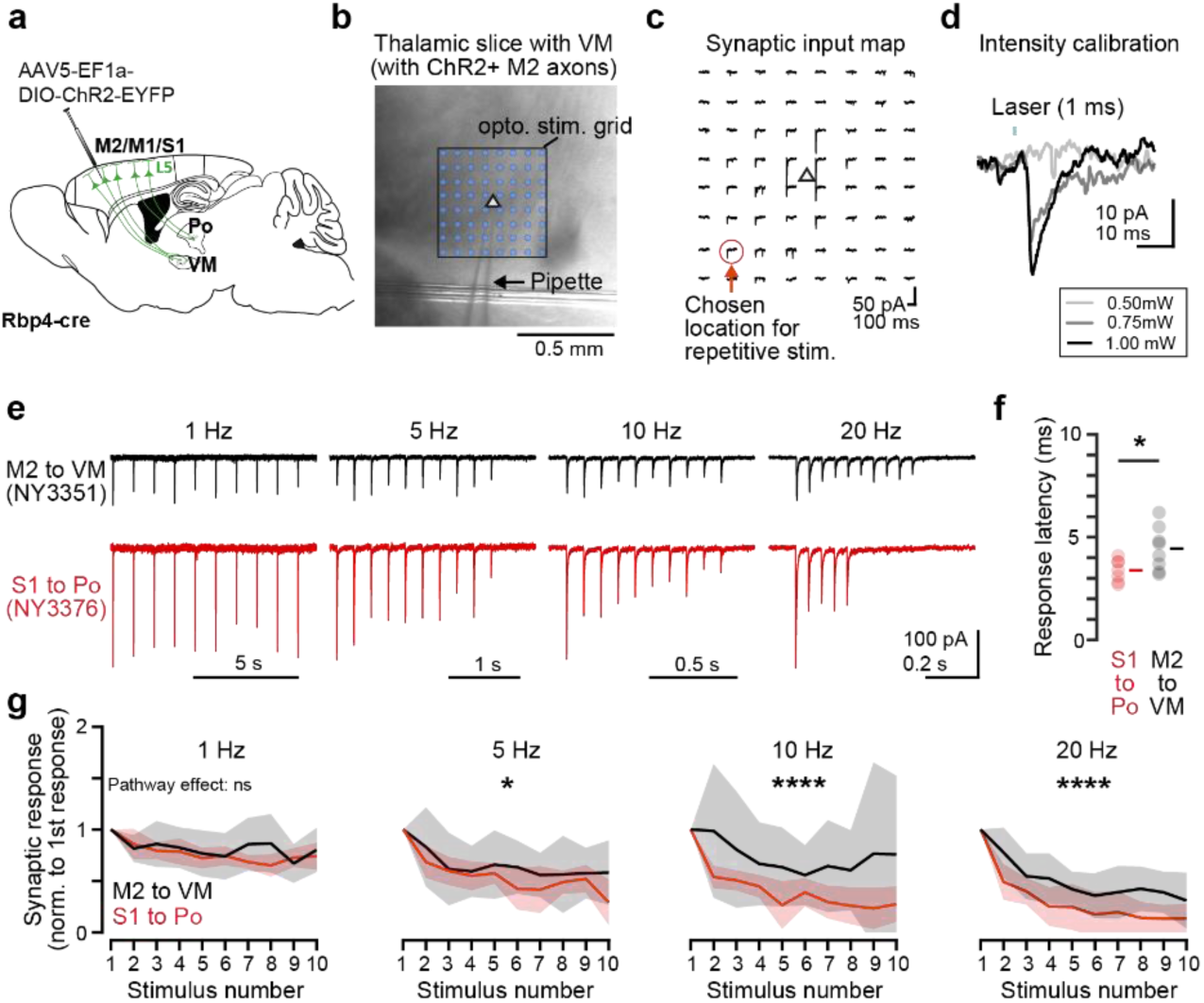
Optogenetic activation of thalamic cells by large S1/L5 and small M2/L5 terminals in vitro. a) Experimental design. b) Representative bright-field image of a thalamic slice with a recording pipette in the VM (arrow) and the overlaid pattern of optogenetic stimulation (delivered for 1 ms at 1 mW). c) Synaptic input map obtained from (b). d) Example traces of photo-evoked EPSCs upon stimulation of the location indicated in (c) at different laser intensities. In this example, 0.75 mW was determined to be used for repetitive stimulation. e) Example traces of synaptic inputs evoked by repetitive stimulation of M2/L5 (black) or S1/L5 (red) axons, recorded from thalamic neurons in VM (NY3351) or Po (NY3376). f) Plot comparing the latency of the first synaptic response to optogenetic stimulation in the power series experiment (see also Extended Data Fig. 5; n = 7 for Po and n = 8 for VM cells). Unpaired T-test, p= 0.035 g) Grouped response amplitude data obtained during repetitive stimulation at each frequency. Synaptic responses to each stimulus were normalized to the first response. Shaded areas represent the SD (n = 6 for Po cells, n = 7 (1, 5Hz) and 8 (10 and 20Hz) for VM cells). Two-way RM ANOVA: main effect of stimulus number: 1 Hz: F(9,110) = 3.1, p = 0.002; 5 Hz: F(9,110) = 5.9, p = 1.1e-6; 10 Hz: F(9,120) = 2.5, p = 0.012; 20 Hz: F(9,120) = 18.9, p = 2.7e-19. Main effect of pathway: 1 Hz: F(1,110) = 2.7, p = 0.103; 5 Hz: F(1,110) = 6.3, p = 0.014; 10 Hz: F(1,120) = 20.0, p = 1.8e-5; 20 Hz: F(1,120) = 32.7, p = 7.9e-5. Interaction between stimulus number and pathway: 1 Hz: F(9,110) = 0.9, p = 0.565; 5 Hz: F(9,110) = 0.5, p = 0.897; 10 Hz:

### Distribution of small L5 terminals throughout the thalamus

As shown above, the small-terminal M2/L5 corticothalamic pathway has a different postsynaptic action from that of the large-terminal S1/L5 pathway, and it selectively targets spines. Thus, we next examined how prevalent the absence of large terminals is among the L5-thalamic pathways arising from frontal cortical areas. We injected AAV-dio-ChR2-EYFP into the M1, M2, frontal associational area (FRA), medial prefrontal (mPFC, including cingulate prelimbic and orbital cortices) insular, S1 and S2 cortices of Rbp4-Cre mice to label L5 pyramidal cells (altogether n=56 animals) (Fig. 6a). Some of these injections have been used before to describe the L5-TRN pathway^40^ (n=29 mice). Injections often included more than one cortical area. Following these injections, large L5 terminals similar to those in Po were found in the n. submedius (Sub), the centrolateral nuclei (CL) and the central part of the mediodorsal nucleus (MDc). In contrast, similar to VM, large L5 terminals (>3 µm) were almost completely missing in the parafascicular n. (PF), paraventricular n. (PV), reuniens n. (Re), rhomboid n. (Rh), lateral and medial MD n. (MDl and MDm), the centriomedial n. (CM), the ventrolateral n. (VL), and ventral anterior n. (VA) (Fig. 6b Extended Data Fig. 6). Cortical regions could send large terminals to one nucleus and small terminals to another (e.g. S1/L5-Po large, S1/L5-PF small. Fig. 6b Extended Data Fig. 6, first column). In order to quantitatively assess the presence or absence of large terminals, we selected 100 AAV-labeled L5 terminals from the high end of the bouton size distribution in the representative sample of 10 L5-thalamic pathways involving the VM, MDl, MDm, SUB, PF, CL and VL nuclei (SUM: 1689 terminals, median: 168/nucleus, range: 115-263/nucleus). In the case of PF and VM we examined L5 inputs from two different cortical sources (M1, S1 and FrA, M2, respectively). We measured the major diameter of L5 terminals in each nucleus using high power light microscopy after DAB-Ni staining and compared them to S1/L5-Po (large L5) or M2/L5-VM (small L5) terminals (Fig. 6b, c). Confirming the qualitative observations, S1/L5 terminals in Po were significantly larger than the L5 terminals in 7 other pathways and were statistically similar only to L5 terminals in mPFC -Sub and FRA-CL (Fig. 6b,c) pathways. Compared to the M2/L5 terminals in VM, the L5 terminals were significantly larger in the mPFC-Sub, FRA-CL, M1-VL and S1-Po but were not different in the other pathways measured (FRA-VM, S1-PF, mPFC-MDM, M2-MDL, Fig. 6b, c). The M1/L5-PF terminals were even smaller than the M2/L5-VM boutons (Fig. 6b, c).

**Figure 6.**
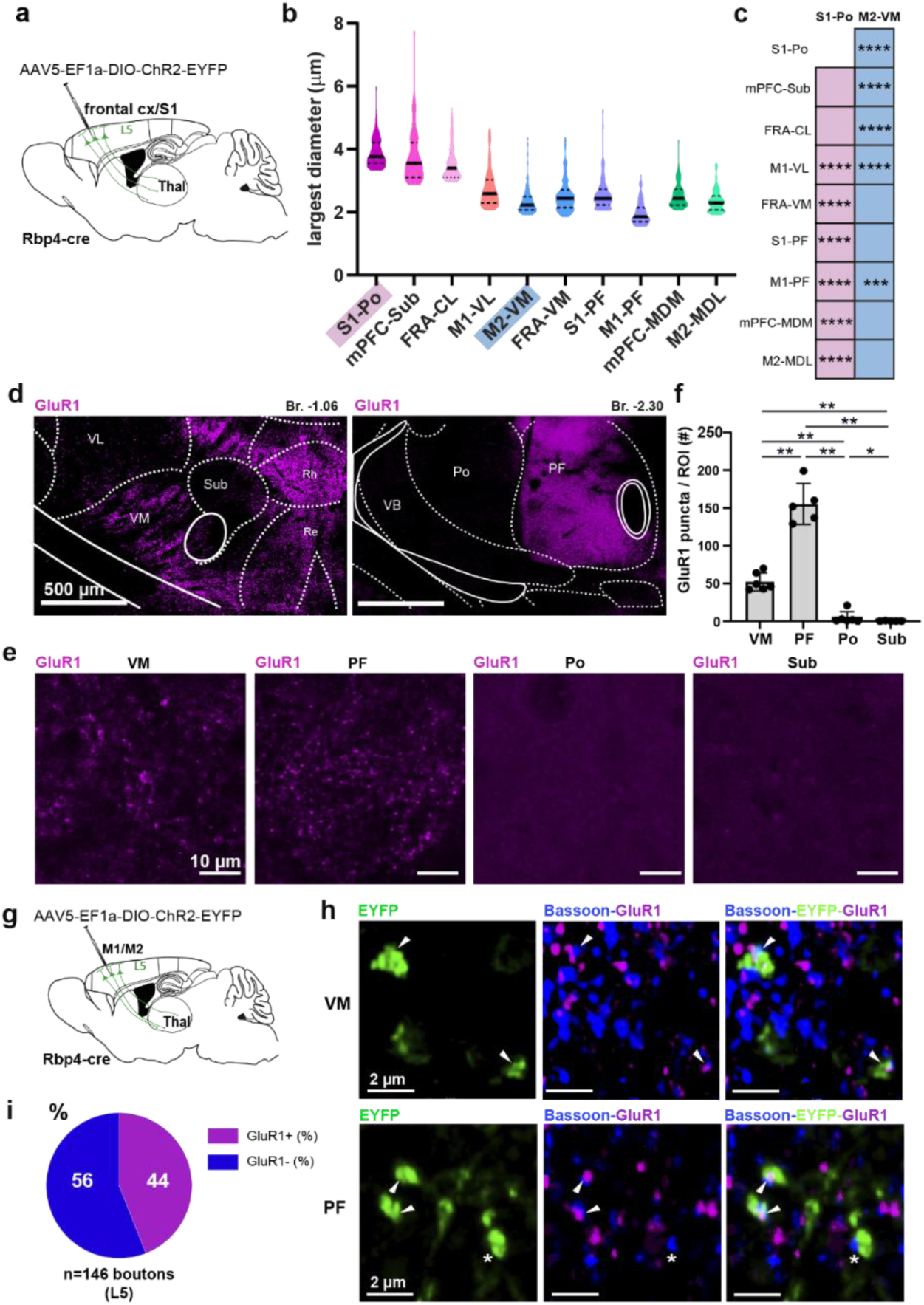
Distribution of small L5-thalamic terminals in the thalamus and their association with GluR1 subunit. a) Experimental design. b) Size distribution of the largest L5 terminals in different corticothalamic pathways. c) Statistical comparison of the terminal sizes in 10 L5 corticothalamic pathways respective to the S1/L5-Po (pink) or the M2/L5-VM boutons (blue). Kruskal-Wallis test with Dunn’s multiple comparison test (all groups were compared to both S1/Po and M2/VM), H(9) = 623,1; p < 0.0001 d) Low power image of immunostaining for GluR1 in two different thalamic regions. e) Representative high power confocal images of GLUR1 immunolabeling from different thalamic nuclei. f) Number of GluR1 puncta per ROI (50 x 50 x 0.755 µm) in nuclei receiving small (VM and Po) or large (Po and Sub) L5 terminals (VM: 52.3±4.8 n=6 mice; PF: 155.4±12.2 n=5 mice; Po: 4.7±3.3 n=6 mice; Sub: 0.2±0.2 n=5 mice; Mann Whitney test: VM-PF: p < 0.01; VM-Po: p < 0.01; VM-Sub: p < 0.01; PF-Po: p < 0.01; PF-Sub: p < 0.01; Po-Sub: p < 0.05). g) Experimental design to label L5 terminals. h) Representative high power confocal images showing synapses (Bassoon, blue) formed by L5 boutons (EYFP, green) facing GluR1 puncta (magenta, white arrowhead). A GluR1 negative L5 synapse is labeled with white asterisk. High power confocal images of EYFP labeled L5 terminals (green), Bassoon (blue) and GluR1 (magenta) in VM (top) or PF (bottom). i) Proportion of GluR1 labeled synapses in the L5-thalamic pathway, sampled from VM and PF.

These data show that L5-thalamic pathways display profound heterogeneity in terminal size. Large thalamic regions including the basal ganglia recipient nuclei (VA, VM, PF), the midline thalamic nuclei (PVT, Re, CM) and parts of the MD receive only small L5 terminals.

### The synapses of small L5 terminals are enriched in GluR1 subunit

Next, we asked whether the small L5-thalamic synapses can contain synaptic proteins that characterize cortical spines as well. We selected the GluR1 subunit of AMPA receptors due to its central importance in cortical spine functions, AMPA mediated synaptic transmission and long-term potentiation^58,59^. Examination of GluR1 immunostained sections at low power demonstrated nucleus specific distributions (Fig. 6d-f), confirming earlier results^50^. Immunopositivity was largely the result of brightly stained immunopositive puncta. Strong immunoreactivity was found only in the nuclei that we found to be contacted exclusively by small L5 terminals (VA-VM, PF, PVT, CM, MDl, MDm, Re) (Fig. 6d-f, Extended Data Fig. 7a), whereas thalamic nuclei innervated by large cortical (SUB, Po, CL, MDc) or subcortical (VPM, VL) terminals displayed weak immunolabeling (Fig. 6d-f, Extended Data Fig. 7a).

To determine whether the bright GluR1-positive clusters are postsynaptic to small L5 terminals, we combined viral tracing in Rbp4-Cre animals from M2 (as above) with immunostaining for GluR1 in the VM and PF (Fig. 6g). We used the presynaptic marker Bassoon to label synapses. Using high power confocal microscopy, we found that in 56% (n=146) of the examined boutons GluR1-positive puncta were located adjacent to the Bassoon signal of small L5 terminals (Fig. 6h,i). As a control we performed the same experiments labeling L6 terminals in Ntsr1-Cre animals (Extended Data Fig. 7b-c). In this case GluR1 immunolabeling could be detected only in 13% of the L6 terminals (n=64 boutons) (Extended Data Fig. 7d,e). These data indicate that the GluR1 subunit is abundant in nuclei receiving only small L5 inputs and synapses of the small L5-thalamic pathways are selectively enriched in the GluR1 subunit.

### Control of thalamic subnetworks by small L5 inputs in freely behaving animals

As shown above, small M2/L5 terminals were able to evoke reliable postsynaptic firing of individual VM cells in anesthetized animals (Fig. 4). To test the efficacy and variability of the response to stimulation of the small L5-thalamic pathway in various thalamic nuclei during different behavioral states, we optogenetically activated their L5 inputs in freely behaving mice. Following the injection of flex-AAV-ChR2-EYFP into the M2 cortex of Rbp4-Cre mice, we implanted an optic fiber and a silicon probe to the M2 cortex and a movable 8-shank silicon probe to the thalamus (n=2 animals). We recorded cortical and thalamic activity during slow wave sleep in the home cage (SWS) and during free exploration (awake) (Fig. 7a-c).

**Figure 7.**
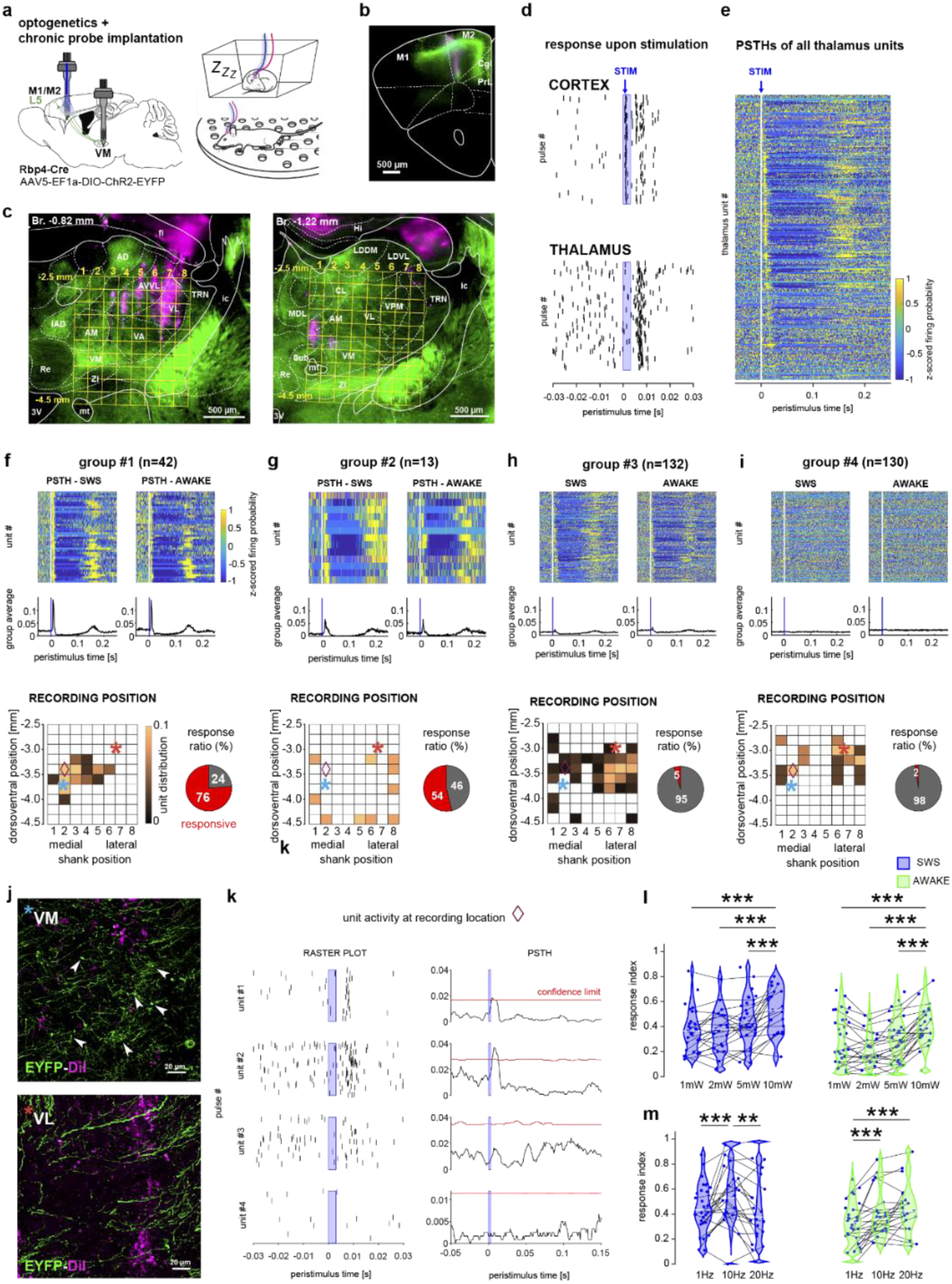
Differential recruitment of thalamocortical cells by small L5 terminals in freely moving animals. a) Experimental design. b) Cortical injection site. Green, anti-EYFP; magenta, DiI, cortical silicon probe track. c) (left) Recording sites in thalamus showing 6 of the 8 DiI labeled shanks (magenta). Yellow grid shows the shank numbers (top) and depth from cortical surface (side). The same coordinates are used in the position maps of f-i panels. (right) The medial- most 2 shanks were located more caudally. d) Representative responses of cortical (top) and thalamic (bottom) units to 1 Hz, 1 mW, 3ms long laser pulse. e) P eri-stimulus time histograms (PSTH) of responses of all thalamic units. Units are organized by the timing of PSTH peak within the first 30 ms after stimulus onset. f-i) Responses of the thalamic units clustered into four types of response groups #1-4. (Top) PSTHs of individual cells in SWS (left) and awake (right) states. (Middle) Population PSTHs for each group. Bottom left: Spatial distribution of the units relative to the recording positions shown in c. Color code shows the proportion of units within a given response group recorded at a given position on the grid. Note the focal concentration of group 1 cells. Bottom right; response ratio calculated as the fraction of units with response higher than the confidence interval (red). Color bar applies for all maps. Asterisks, the locations of images in j and k; diamond, the location of recordings shown in k. j) (Upper panel) High power confocal images of ChR2 labeled fibers (green) with small L5 axon terminals (arrowheads) around recording positions (magenta, DiI labeling of shank 2) where Group 1 neurons were frequent (shank 2, blue asterisk in f,). (Lower panel) ChR2-positive axons with only passing fibers around recording positions where mainly non-responsive Group 4 neurons were recorded (shank 6). k) Representative evoked responses of four different units with different response characteristics recorded within 150 μm in shank 2 (purple diamond in f). l-m) Response probability indices of the significantly activated group #1 units by increasing illumination power (l) or using different stimulation frequencies (m), in both SWS and awake states. (l) Two-way RM ANOVA results: intensity effect: F(3,60) = 39.978, p = 2.5283×10^−14^, state effect: F(1,20) = 24.353, p = 7.9837×10^−5^, interaction effect: F(3,60) = 1.9746, p = 0.12738; asterisks indicate intensity effects within each state (post-hoc comparison by Tukey’s HSD test). (m) Two-way RM-ANOVA results: frequency effect: F(1,21) = 19.191, p = 1.2023×10^−6^, state effect: F(1,21) = 11.299, p = 0.0029534, interaction effect: F(2,42), p = 0.00091827; asterisks indicate frequency effects within each state (post-hoc comparison by Tukey’s HSD test).

At the injection site, cortical ChR2-positive L5 cells (n=20) responded to the laser stimulation (1 mW, 1 Hz) with short latency (median latency: 5 ms) and high fidelity (median response index: 0.469) (Fig. 7d, Extended Data Fig. 8a-c). In the thalamus many units were also activated by L5 stimulation, but their peristimulus time histogram (PSTH) displayed large heterogeneity (Fig. 7d-e). The PSTHs of thalamic units were organized by heuristic K-means clustering into 4 groups with different response characteristics (Fig. 7f-i). The firing activity of most (76%) Group 1 neurons (n=42) were significantly modulated by L5 stimulation (i.e., exceeded the 95th percentile of the randomly generated PSTHs obtained through a surrogate test (95% Fig. 7f). In this group the response onset was fast (peak median latency: 7 ms) followed by a decrease in firing and a rebound at ∼150 ms. *Post hoc* anatomical analysis revealed that these units were localized in the VA, VM nuclei and the electrode tracks labeled by DiI at these depths were surrounded by ChR2-EYFP-positive small L5 terminals (Fig. 7c, j upper panel). Group 2 cells (n=13) had significantly slower response onset than Group 1 neurons (median: 12 ms, p<0.001, Mann Whitney U test). These cells were located outside the thalamus; thus, they were not considered here for further analysis (Fig. 7g). The PSTHs of Group 3 cells (n=132) displayed a small initial activation followed by a decrease in firing and a rebound at ∼150 ms (Fig. 7h). Only 5 of the Group 3 cells (3.78 %) displayed significant response to cortical stimulation. Finally, Group 4 cells (n=130) displayed no modulation to cortical stimulations (Fig. 7i). Group 3 and 4 cells were mainly located in VL, where only passing ChR2-EYFP fibers were observed (Fig. 7j, lower panel). The population PSTHs of the four groups did not change when the same neurons were recorded during active exploration (Fig. 7f-i).

Unresponsive or weakly responsive Group 3 and 4 neurons were also located in VA/VM intermingled with Group 1 cells (Fig. 7k) showing that thalamic neurons with distinct response probabilities could be recorded in close vicinity to each other. At the population level this resulted in highly heterogeneous mean response probabilities even in neighboring recording positions within the focus of the L5 termination field (Extended Data Fig. 8d, shank 2-4, dv. positions 3.0-3.6 mm). Response probability did not depend on the spontaneous firing rates (Extended Data Fig. 8e) indicating that it is not simply the result of higher excitability. The response probability of significantly activated Group 1 thalamic units displayed great heterogeneity in both the SWS and the awake states (Fig. 7l).

Increasing the intensity of the cortical stimulation from 1 to 10 mW increased cortical response probabilities (Extended Data Fig. 8c) as was also shown before^40^. Despite the stronger L5 excitation the response probabilities of Group 1 thalamic units did not change significantly to 1 mW, 2mW and 5 mW photoactivation, indicating that increasing the number of activated L5 cells does not necessarily lead to stronger firing output (Fig. 7l). Thalamic cells started to respond with higher probability only when 10 mW laser power was applied. When we compared the two behavioral states the mean response probabilities of activated Group 1 neurons were significantly smaller in the awake states compared to sleep at all laser powers (Fig. 7l, Extended Data Fig 8g, group 1).

Contrary to the in vitro and anesthetized conditions, increasing the frequency of M2/L5 stimulation did not result in significant decrease in the response probabilities in Group 1 cells in both awake and sleep (Fig. 7m). The response latency remained fast in every condition and did not display state dependent changes (Extended Data Fig. 8f).

These data show that in freely behaving conditions small L5-thalamic inputs can efficiently recruit thalamic cells with pronounced cell-to-cell variability even among neighboring neurons. Recruitment was state dependent, showed facilitation at higher frequencies but was less influenced by increasing activation power.

### Perturbation of small L5-thalamic synapses impairs motor learning

M2 cortex is known to be involved in motor learning and representing the 3D posture of the animals^41,42^. Thus, we finally tested if interference with the small M2/L5-thalamic communication could affect the acquisition of a novel motor skill (Fig. 8a). We selected a spontaneous horizontal wheel learning paradigm since it does not include the potentially confounding effects of punishment or reward and can be easily quantified. Well-handled but task-naïve mice were exposed to a running wheel for one hour per day for seven days while they spontaneously acquired the wheel running task. We used open field and place preference tests to control for the specificity of the motor learning. Rbp4-Cre mice were injected in the M2 with AAV5—flex-ArChT-tdTomato or control virus AAV5-CAG-FLEX-tdTomato. Optic fibers were implanted bilaterally into the VM (Fig. 8b). The laser light was activated in a closed-loop manner when the animals entered the wheel area, using a 5 s ON/ 10 s OFF duty cycle that has been successful previously to perturb L5 activity in the TRN^40^. Following histological verification (Fig. 8b) we compared the performance of the Ctrl (n= 9) and ArChT (n= 9) animals during the initial (1-3 days) and late (5-7 days) phase of the learning (Fig 8c).

**Figure 8.**
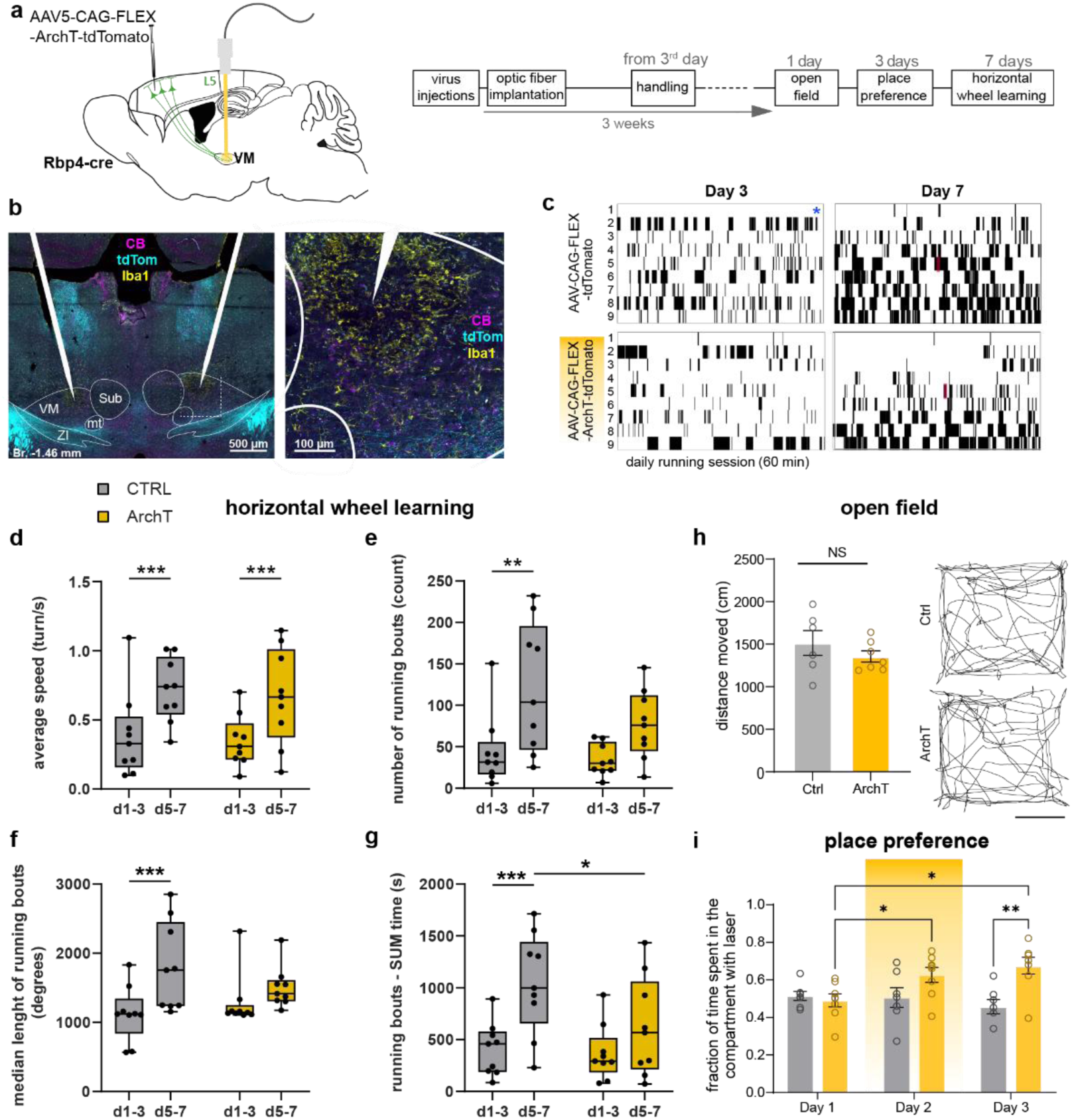
Optogenetic perturbation of the M2/L5- VM pathway interferes with the acquisition of a motor learning task. a) Schematic overview of surgery (left) and experimental procedure (right). b) Confocal images of optic fiber placement and L5 axon arbor (tdTom, blue) in the VM. Iba1, (yellow) microglia activation to label optic fiber position, CB, (blue) calbindin. c) Rasterplots of running bouts (min 3 turns, black rasters) for each animal at day 3 and 7 (Ctrl top, ArchT bottom). Individual rows depict individual animals. Blue star, the animal did not reach running criteria, red boxes indicate the time of supplementary videos (see Extended Data). d) Average speed of animals over all running sessions (day-1-3 compared to day 5-7) (2-way repeated measures ANOVA followed by uncorrected Fisher’s LSD, interaction: F(1, 16) = 3.971×10^−3^, P=0.9505, group: F (1, 16) = 0.1322, P=0.7209, time: F (1, 16) = 42.25, P < 0.0001, e) Average number of running bouts (day-1-3 compared to day 5-7) (2-way repeated measures ANOVA followed by uncorrected Fisher’s LSD, group: F(1, 16) = 2.771, P=0.1154, time: F(1, 16) = 11.97, P=0.0032, interaction: F(1, 16) = 0.9436, P=0.3458). f) Median length of running sessions (day-1-3 compared to Day 5-7) (2-way repeated measures ANOVA followed by uncorrected Fisher’s LSD, group: F(1, 16) = 0.1177, P=0.7360, time: F (1, 16) = 18.32, P=0.0006, interaction: F(1, 16) = 4.860, P=0.0425). g) SUM time of running bouts, (day 1-3 compared to day 5-7) (2-way repeated measures ANOVA followed by uncorrected Fisher’s LSD, group: F(1, 16) = 2.276, P=0.1509, time: F(1, 16) = 19.90, P=0.0004, interaction: F(1, 16) = 3.861, P=0.0670)). h) Distance moved by animals in the open field during a 3-minute period receiving laser stimulation (unpaired t-test, p=0.3218). Scale bar, 10 cm. Right, trajectory of two representative animals in the open field.

Both control and ArchT mice showed improvement in performance with experience, such as an increase in average speed and in the number of running bouts (Fig. 8d, e), indicating that motor activity and motivation of the animals were not affected by the optogenetic perturbation. However, compared to control animals, the length of running bouts increased significantly less in ArchT mice (Fig. 8f) and accordingly they spent overall less time running in the later sessions (Fig. 8g).

We could also exclude any gross disruption of motor functions since we found no significant difference between ArchT (n=7) and control (n=6) animals in the open field test (Fig. 8h). We could also exclude any aversive effect of the ArchT activation in VM using the place preference test (Fig. 8i). In fact, *post hoc* analysis of the ArchT group (n=8) showed significant increase of time spent in the stimulated compartment, an effect which persisted on the third non-stimulated day (Fig. 8i).

In sum, we conclude that closed-loop perturbation of small L5-thalamic connections prevented the improvement in several measures in a motor learning paradigm. The results could not be explained by a general deficit in motor control or by a conditional place aversion induced by the optogenetic inhibition but suggests a deficit in the acquisition or consolidation of advanced motor skills.

Place preference task. Fraction of time spent in the compartment in which animals received laser. Two-way ANOVA, followed by Sidak’s multiple comparisons test. groups (F=5.975, p=0.029) interaction (F=7.109, p=0.0034). *Post hoc* analysis of experimental groups showed significant increase of time spent in the compartment on day 3 in ArchT (n=8) animals compared to the control (n=7) group (p=0.0026). *Post hoc* analysis of temporal changes in experimental groups showed increased time spent in the compartment in the ArchT group between day 1 and day 2 (p=0.0302) and day 1 and day 3 (p=0.0294)

## Discussion

Our data disclosed a novel type of communication channel between the cortex and the thalamus that involves specialized contacts between L5 terminals originating in the frontal cortex and functional thalamic spines. To the best of our knowledge, this is the first demonstration of specific axo-spinous signal transfer by a pathway targeting excitatory cells in a subcortical structure.

The axo-spinous contacts established by small L5 corticothalamic terminals are distinct from the previously described corticothalamic pathways (large L5 and small L6 pathways) both in morphology and physiology^24,60^. Small L5 terminals differ from large corticothalamic L5 terminals in that they target thalamic spines via a single postsynaptic element, not complex dendritic appendages with multiple synapses^15,17,18^. The spines postsynaptic to small L5 terminals contain spine apparatus unlike the dendritic appendages postsynaptic to large L5 terminals^26^. Small L5 terminals differ from small L6 terminals in that the latter contact dendritic shafts, not spines, and unlike L6 boutons small L5 terminals contain mitochondria. Large L5 terminals are considered as “driver” inputs in the thalamus and display strong short-term depression, whereas small L6 terminals are considered as “modulatory” input and display short-term facilitation^24,54,60^. The postsynaptic action of small L5 corticothalamic terminals described here differed from both. They could drive postsynaptic firing with high (but variable) probability even at high input frequencies. Absence of large L5 corticothalamic terminals characterized many thalamic regions receiving frontal cortical inputs indicating that this mode of information transfer is abundant in corticothalamic pathways responsible for motor and executive functions.

To our knowledge up till now no specific afferents have been associated with classical dendritic spines in the thalamus. The comparison of small L5-thalamic and cortico-cortical axo-spinous contacts reveals several common features. Both have comparable spine head and neck dimensions and can contain spine apparatus^4,19,47^. Spine head and PSD dimensions strongly correlate in both cortical and thalamic spines^47^. Spines in both regions are mainly contacted by single terminals via one synapse which can have complex shape (perforated, branching)^19^ and can be enriched in GluR1 subunit^59,61^. Finally, we demonstrate here that thalamic spines display compartmentalized Ca^2+^ responses similar to cortical (i.e., hippocampal) spines.

Spines are considered to be the postsynaptic substrate of plastic synaptic alterations in cortical neurons, where increase or decrease in synaptic strength are associated with changes in spine size and shape^8,62,63^. Increased spine size is thought to accompany a rapid increase in synaptic transmission by enhanced AMPA receptor insertion into the postsynaptic membrane upon long-term potentiation^8,63^. The presence of GluR1 subunit in the AMPA receptor complex was proposed to facilitate synaptic plasticity and compartmentalized Ca^2+^ responses are considered to enable input-specific activity-dependent modification of synaptic strength^58,61,64^. Our finding that the small cortical L5 axons preferentially target spines that express GluR1 subunit raises the possibility that these synapses may also undergo synaptic plasticity. The exact conditions of thalamic synaptic plasticity - if present – may, however, follow different rules, since the dendritic properties, connectivity and spontaneous activity of pyramidal cells are substantially distinct from thalamocortical cells.

In freely behaving conditions the impact of small L5 terminals from the frontal cortex on thalamocortical cells was characterized by three features. 1. Optogenetic activation of small L5 inputs resulted in significant but highly variable thalamic responses. 2. Stable output could be generated using variable stimulation frequencies and power. 3. The response properties were state dependent. In theory, these features could allow stable representation of a cortical input pattern by a well-defined thalamocortical cell population even during changing input conditions. Since thalamic territories innervated by small L5 terminals project to widespread cortical and subcortical regions^65^, communication via this “trans-thalamic” route could have large, differentiated impact on forebrain networks.

Our present data can provide insights into the mechanisms behind these three features. Small L5-thalamic synapses could evoke postsynaptic firing with high, albeit variable, efficacy, as even the smallest stimulation intensities (0.01 mW) could activate some of these cells with relatively high probability. We attribute the effectiveness of the small L5-thalamic contact to both pre- and postsynaptic morphological features. The presynaptic small L5 terminals almost always contained mitochondria, unlike L6 terminals and many cortical terminals^66^. The presence of mitochondria indicates high metabolic capacity in these boutons^67^. In addition, the proximity of a mitochondrion has a positive influence on synaptic performance^68^. Accordingly, we found here that the small L5 terminals - which always had mitochondria - established significantly larger synapses than the L6 boutons lacking this organelle. On the postsynaptic side the PSD surface area of small L5 terminals was highly variable, some of them were quite large. Indeed, at the higher end of this spectrum the individual small L5 synapses were comparable in size to the combined PSD surface of the many, small, macular synapses established by a single large thalamic “driver” terminal^17,18^. Since the PSD area scales with AMPA receptor content^69^, charge transfer through these L5-thalamic synapses can evoke strong postsynaptic response. The response probabilities of thalamic cells were not uniformly high to L5 stimulation but displayed large cell-to-cell variability: non-responsive and moderately to highly responsive neurons were intermingled even in the thalamic hot-spots of the L5-thalamic pathway. Variable synaptic strength at the L5 axo-spinous synapse is probably an important factor in this heterogeneity, since the L5 release sites spanned tenfold difference in size at the EM level. In addition, the variable number of L5 inputs that any given thalamic neuron receives may also be important. Curiously, increasing the number of recruited L5 cells by increasing stimulation power in L5 between 1 and 5 mW^40^ did not result in increased thalamic response probabilities at the population level, instead the output remained stable. Interestingly, using a very similar approach to stimulate the L5-TRN pathway, the response in the TRN scaled precisely with the increasing stimulation power^40^. In fact, this L5-TRN connection may participate in the normalization of thalamic output during variable input conditions via feed-forward inhibition, since the TRN sectors activated by L5, target the same thalamic nuclei, that are innervated by small L5^40^ terminals.

Small L5-thalamic pathway displayed STD *in vitro*. In the freely behaving condition, however, response probabilities remained stable at every stimulation frequency tested indicating that in the more natural condition these thalamocortical cells can reliably read out the input frequency. In an earlier study^39^, the thalamic nuclei examined here displayed ramping activity similar to that of their cortical input cells during the delay period of a sensory-motor working memory task. This thalamic activity pattern was sensitive to cortical inactivation^39^. Effective transfer of L5 information by thalamic neurons via the small cortical L5 terminals may require stable output at different frequencies, as we found here. In the above sensory-motor task, cortical inactivation impaired the performance of the animals. In our case, perturbation of only the L5 axonal activity in the VM was sufficient to impair the acquisition of a motor skill that requires coordinated activity of several motor centers and in general a highly precise ensemble activity of motor thalamus^70^. A recent study^36^ observed STD in the thalamic response for 20 Hz stimulation of L5 whereas in our case the thalamic output remained stable at 20 Hz. Different experimental conditions (e.g. brief stimulus train vs steady state responses following longer trains, head-restrained vs freely moving condition) can explain these differences.

Thalamic responses were significantly stronger during SWS compared to wake state at every condition tested. This is somewhat counterintuitive but may be explained by state-dependent differences in excitability of thalamocortical cells, which are endowed with highly nonlinear active conductances^71,72^. High sensitivity of thalamic responses to cortical inputs may allow the small L5-thalamic network to participate in sleep replay.

We studied the ultrastructure of the small L5 terminals and their postsynaptic partners only in VM. However, the presence of small L5 terminals and its association with GluR1 subunits in other nuclei strongly suggest that L5 axo-spinous contacts are more widespread in the thalamus. Small L5 terminals have also been found in the Po^45^ (and this study) but their postsynaptic targets remain to be established. Furthermore, using Golgi staining and intracellular filling, spines have been observed in the first order sensory thalamic nuclei that are known to have no layer 5 inputs^12^. If those spines also have synaptic inputs, axo-spinous contacts may not only be formed by small L5 afferents in the thalamus. However, the paucity of GluR1 immunostaining in Po and the other nuclei contacted by large excitatory terminals indicate that in these regions, even if axo-spinous contacts are present, their synaptic properties probably differ from those of the small L5 contacted spines.

Axo-spinous communication among excitatory cells has been dominantly attributed to intracortical networks. Here we describe a region-specific, top-down, corticothalamic pathway arising primarily from the L5 of frontal cortices that target functional spines in excitatory thalamocortical cells. These connections can provide a versatile form of forebrain communication via thalamic nuclei with widespread connections. Our results have the potential to expand research on novel functional roles of thalamocortical circuits.

## Acknowledgements

We thank the Light Microscopy Center, the Electron Microscopy Center and the Virus Technology Unit at the Institute of Experimental Medicine for providing support for the experiments. Authors would like to thank Krisztina Faddi, Zsófia Varga-Németh, Szabina Seres for their excellent technical assistance, Noémi Kis for her help in the initial phase of the in vitro calcium imaging experiments and Seba Aljoomaa for her help in bouton size quantification. LA was funded by the ERC Advanced Grant FRONTHAL, 742595 (LA) the European Union project within the framework of the Artificial Intelligence National Laboratory RRF-2.3.1-21-2022-00004 and the “Lendület” Program of the Hungarian Academy of Sciences (grant no. LP2023-2/2023), GS was funded by NIH/NINDS R01/R37 and the Javits Award NS061963. JKM was supported by the European Research Council (CoG 771849), the International Research Scholar Program of the Howard Hughes Medical Institute (55008740), the National Research, Development and Innovation Office, Hungary (Highlight 152205) and the National Brain Research Program (NAP3.0) of the Hungarian Academy of Sciences (NAP2022-I-1/2022). This research work was conducted with the support of the National Academy of Scientist Education Program of the National Biomedical Foundation (BT).

## Extended Data Figures

**Extended Data Fig. 1.**
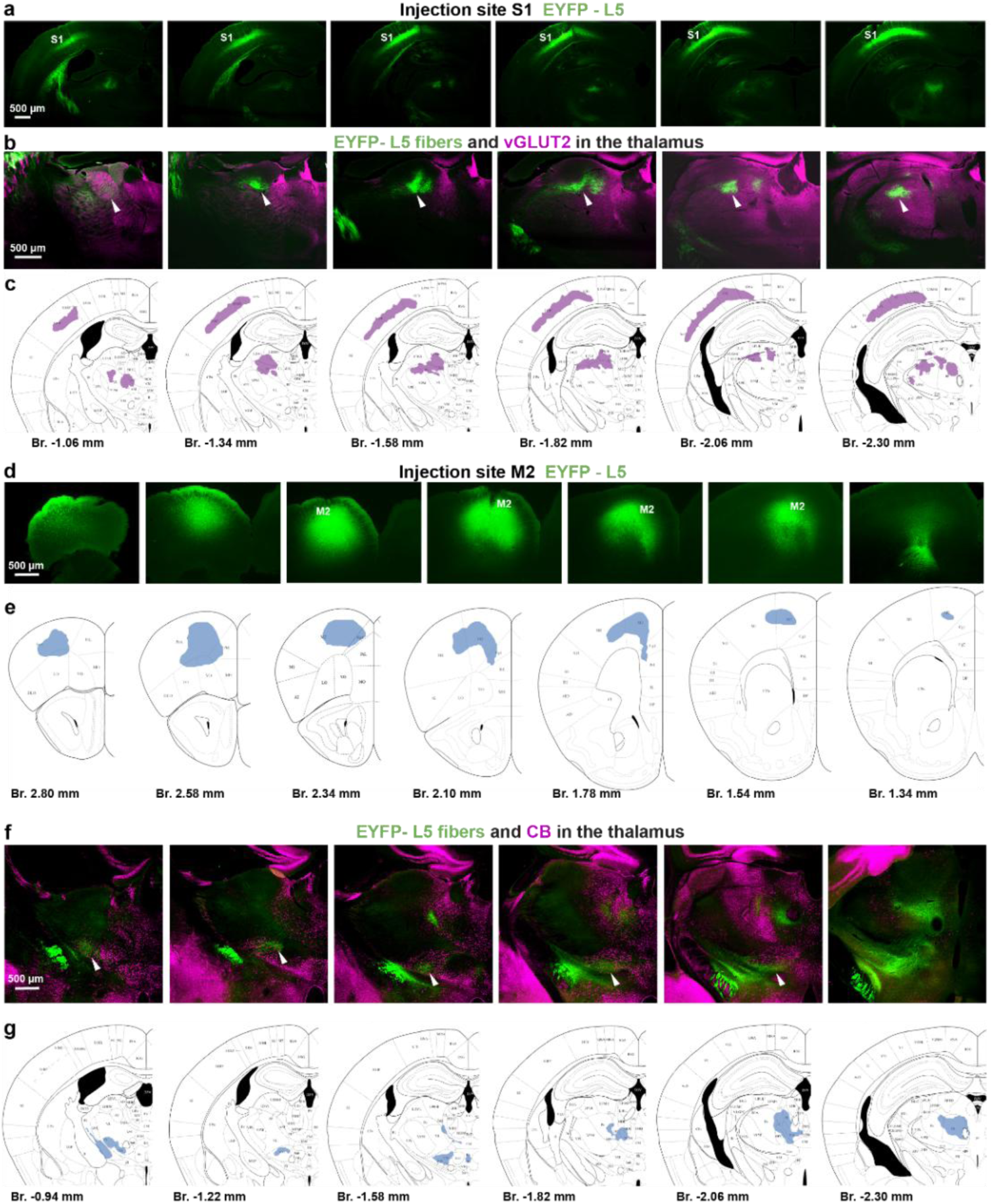
Distribution of L5 fibers in the thalamus following S1 and M2 injections. Related to Figure 1. a) Cortical injection site of a representative S1/L5 injection at six antero-posterior levels (EYFP, green). b) Termination field of L5 fibers (EYFP, green) in the thalamus, sections immunostained also for vesicular glutamate transporter-type-2 (vGLUT2, magenta). c) Schematic drawing of the injection site and the distribution of thalamic fibers. d-e) A representative M2/L5 injection site (EYFP, green). f-g) Termination field of M2/L5 fibers in the thalamus, sections immunostained also for calbindin (CB, magenta).

**Extended Data Fig. 2.**
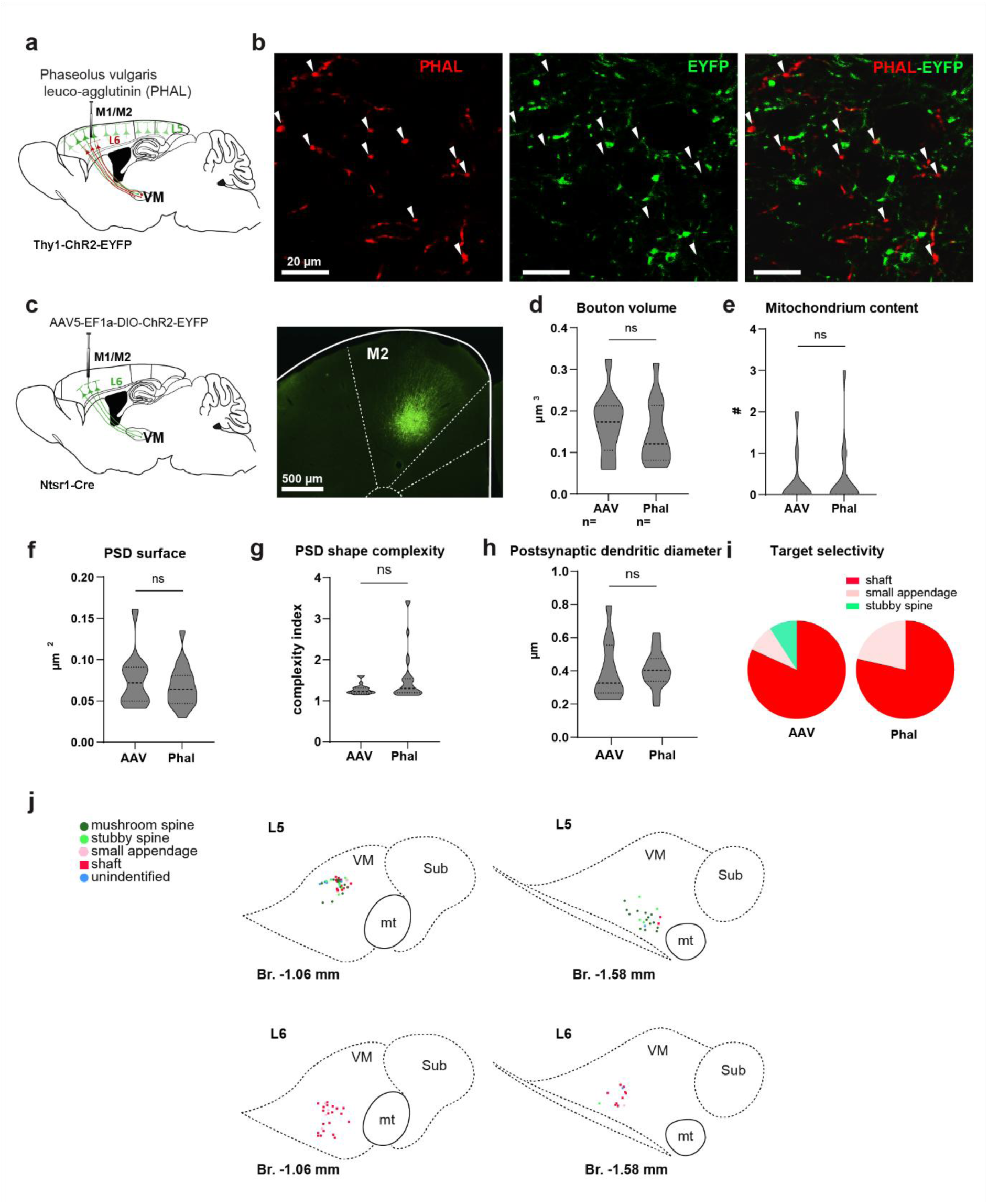
Properties of AAV and PHAL labeled L6 boutons and position of L5 and L6 boutons analyzed in Figure 2. Related to Figure 2. a) Experimental design for labeling L6 cells with PHAL in the Thy1-EYFP mouse, expressing the EYFP-ChR2 fusion protein (green) in L5 cells and terminals. b) Thy1-L5 (EYFP, green) and PHAL-labeled (red) axon terminals in the VM. Left and middle: single channels, right: overlay. Note that the two populations do not overlap, showing that the PHAL injection targeted the L6 cells. c) Left: Experimental design for viral (AAV, left) labeling of L6 cells. Right: injection site. d) Bouton volume (AAV: 0.17±0.02 µm^3^ n=11; PHAL: 0.15±0.02 µm^3^ n=13; Mann Whitney test: ns) e) Number of mitochondria (AAV: 0.27±0.19 n =11; PHAL: 0.31±0.20 n=16; Mann Whitney test: ns) f) PSD surface (AAV: 0.08±0.01 n =11; PHAL: 0.07±0.01 n=17; Mann Whitney test: ns) g) PSD shape complexity (AAV: 1.29±0.04 n =11; PHAL: 1.51±0.13 n=20; Mann Whitney test: ns) h) Postsynaptic dendritic diameter (AAV: 0.40±0.05 n =11; PHAL: 0.41±0.02 n=26; Mann Whitney test: ns) i) Postsynaptic element type (AAV: shaft 81.8% small appendage 9.1% stubby spine 9.1% mushroom spine 0% n=11; PHAL: shaft 78.57% small appendage: 21.43% stubby spine: 0% mushroom spine: 0%, n=28) j) Position of the boutons analyzed in Fig. 2 within the VM. Boutons are labeled with different colors according to their postsynaptic target (dark green: mushroom spine; light green: stubby spine; pink: small appendage; blue: type of postsynaptic target could not be identified. Violin plots: horizontal endpoints, minimum and maximum values; scattered lines, first to third quartiles; thick dashed line: median.

**Extended Data Fig. 3.**
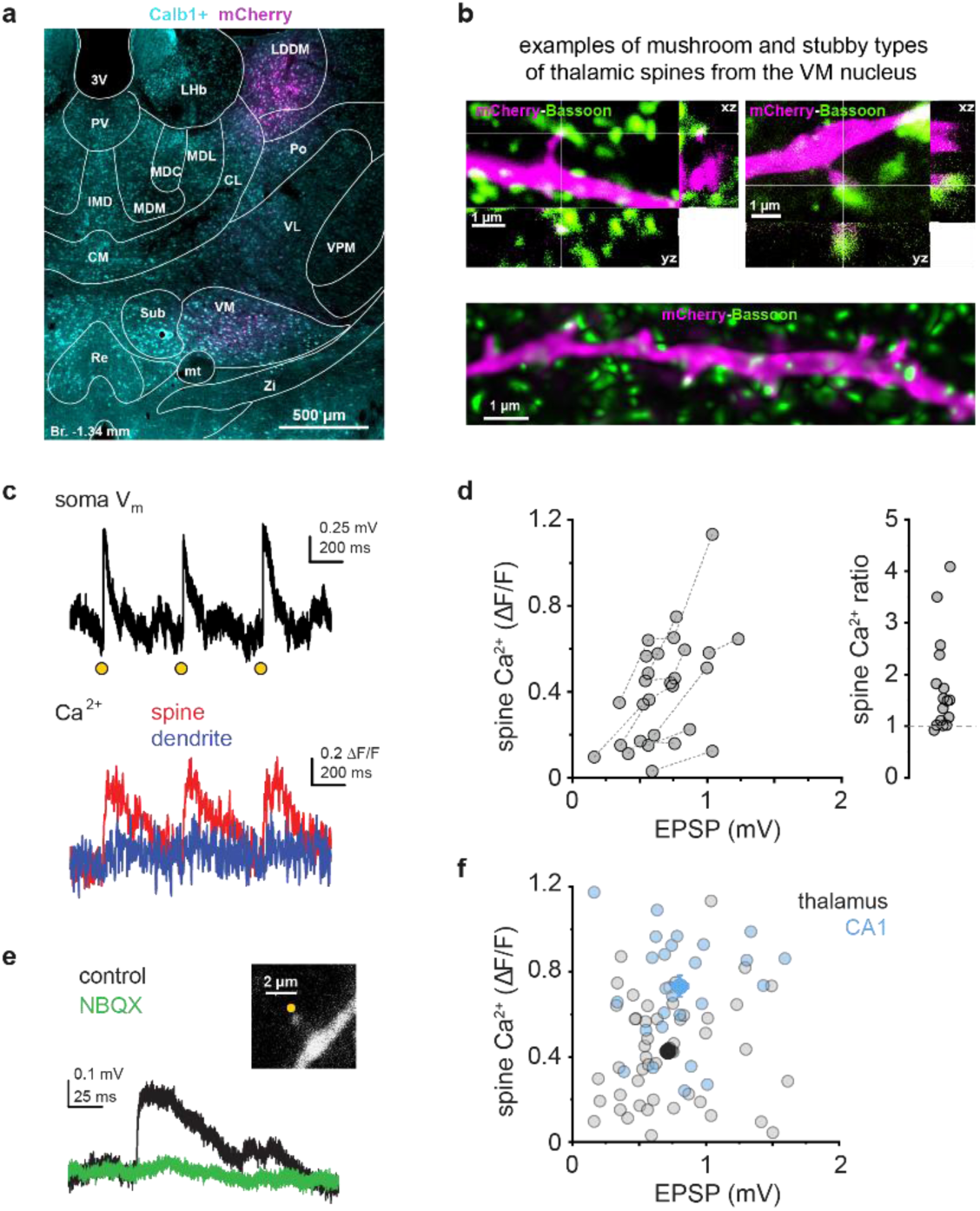
Details of 2P Ca^2+^ imaging experiments. Related to Figure 3. a) Low power image of an injection site. b) Examples of thalamic spines with different morphology. (Top) maximum intensity projections of 3D STED images (x, y, z: 30 nm pixel size) of thalamic dendritic segments bearing a mushroom spine in close contact with a terminal labeled by the presynaptic protein Bassoon. (Bottom) deconvolved confocal reconstruction of a longer thalamic dendritic segment with stubby spines, some of which receive putative contacts labeled with Bassoon protein. c) Response of a thalamic spine to repetitive 2PGU. (Top) somatic voltage response; (bottom) average spine and dendrite Ca^2+^ response to 3 repetitions of 2PGU (yellow dot) within a trial. d) Left, 2PGU-evoked EPSPs and corresponding spine Ca^2+^ signals in spines that were stimulated in multiple 2PGU series using different stimulation settings (laser power, duration and/or position around the spine head), producing EPSP pairs with > 10% different amplitudes and <1.3 mV amplitude. Right, ratio of spine Ca^2+^ signals in individual spines (Ca^2+^_Larger EPSP_/Ca^2+^_Smaller EPSP_, mean ± SEM: 1.77 ± 0.23, n=16, Wilcoxon test comparison to 1: p<0.001). e) Representative average voltage responses to 2PGU in a thalamic spine (inset, single scan) before (black) and 10 minutes after (green) bath application of 20 μM NBQX. Similar effect by 10 μM NBQX was observed in two other thalamic neurons. f) 2PGU-evoked EPSPs and corresponding spine Ca^2+^ signals in CA1PC spines (blue dots, n=27 stimulation series in n=21 spines in 5 cells) and thalamic spines (grey dots, n=49 stimulation series in n=27 spines in 18 cells, same data as in Fig. 3j, m). Large filled symbols: mean ± SEM of data measured in the two cell types. EPSP amplitudes were similar (Mann-Whitney test, p=0.125) whereas corresponding spine Ca^2+^ signals were larger in CA1PCs (Mann-Whitney test, p<0.001).

**Extended Data Fig. 4.**
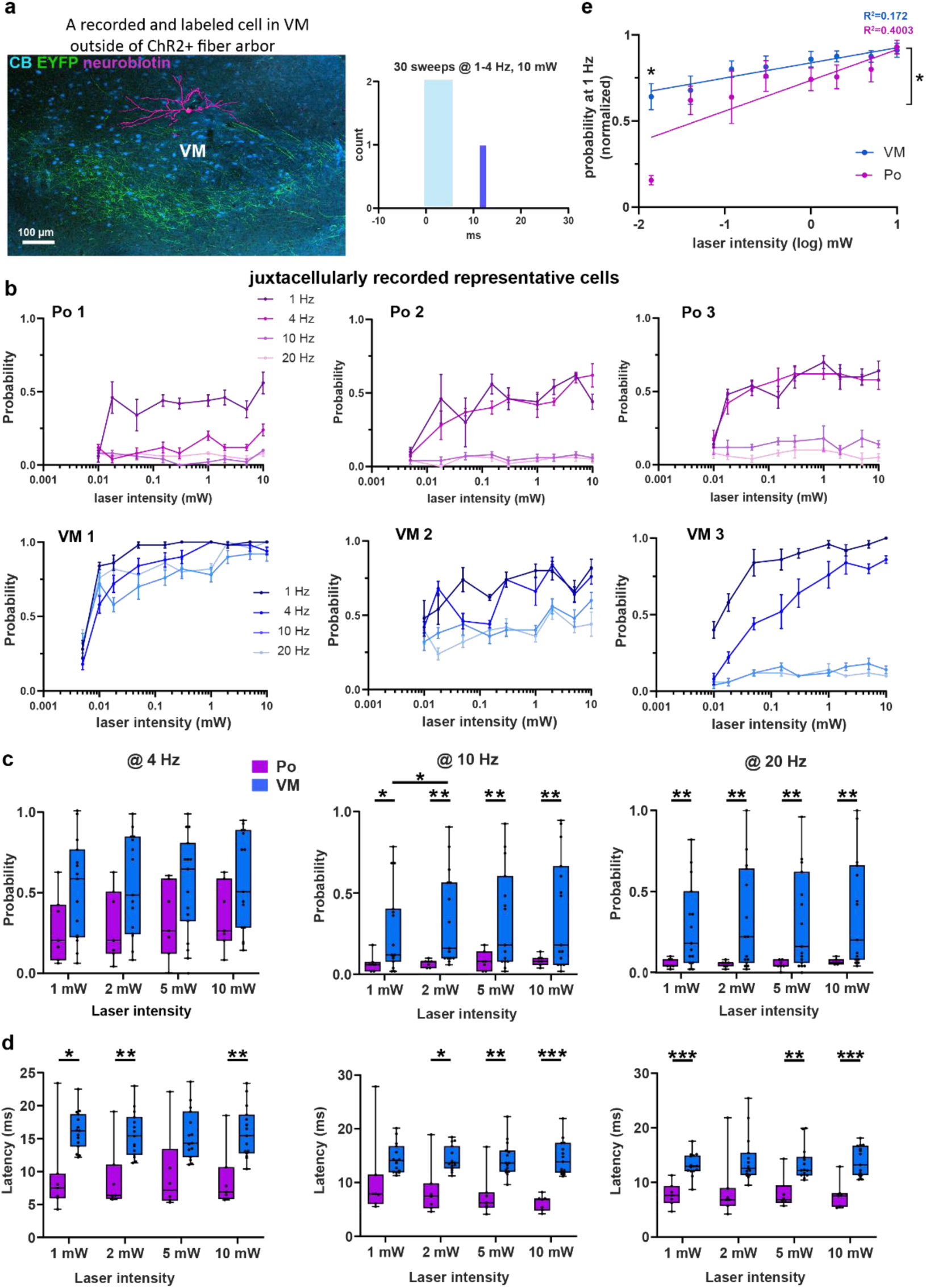
Details of the impact of optogenetic L5 stimulation on VM and Po cells. Related to Figure 4. a) Example of a recorded and juxtacellularly labeled VM cell outside of the ChR2 positive axons form M2 L5 cells. PSTH in the middle panel shows that 10mW stimulation of ChR2+ cells in M2 could not elicit responses from this cell. b) Response probabilities of individual representative cells from the Po (Po-1-3) and VM (VM 1-3). Responses for 1, 4, 10, 20 Hz cortical (mean ± SEM, S1 for Po, M2 for VM cells respectively) stimulation with increasing laser intensity are shown for each cell. c) Response probability of the Po and VM population for 4, 10 and 20 Hz stimulation at increasing laser intensities. (Two-way 2-way RM ANOVA with GG correction, followed by Tukey’s *post hoc* tests for all. *4 Hz (*as none of the ANOVA tests were significant, no *post hoc* test is shown): intensity: F(3,60) = 1.284, GG ε = 0.5748, p=0.2862, nucleus: F (1,20) = 3.695, p=0.0689,, interaction: F(3,60) = 0.08061, GG ε = 0.5748, p=0.8981. *10 Hz*: intensity: F(3,60) = 2.584, GG ε = 0.5020, p= 0.1047, nucleus: F (1,20) = 5.182, p= 0.0340, interaction: F(3,60) = 1.380, GG ε = 0.5020, p= 0.2623; *20 Hz*: intensity: F(3,60) = 1.334, GG ε = 0.5446, p= 0.2733, nucleus: F (1,20) = 5.483, p= 0.0297, interaction: F(3,60) = 0.8083, GG ε = 0.5446, p= 0.4318. d) Response latencies of the Po and VM population for 4, 10 and 20 Hz stimulation at increasing laser intensities (4 Hz: mixed-effect analysis with GG correction, followed by Tukey’s *post hoc* tests: intensity: F(3,60) = 0.2188, p = 0.8043, GG ε = 0.6661, nucleus: F (1,20) = 14.32, p = 0.012, interaction: F(3,60) = 0.09112, GG ε = 0.6661 p = 0.9130. *10 Hz*: 2-way repeated measures ANOVA with GG correction, followed by Tukey’s *post hoc* tests: intensity: F(3,60) = 3.020, p= 0.0701, GG ε = 0.5630, nucleus: F (1,20) = 18.72, p= 0.0003, interaction: F(3,60) = 3.188, p= 0.0616, GG ε = 0.5630. *20 Hz*: mixed-effect analysis with GG correction, followed by Tukey’s *post hoc* tests: intensity: F(3,60) = 0.5426, p = 0.5333, GG ε = 0.4891, nucleus: F (1,20) = 25.52, p < 0.0001, interaction: F(3,60) = 0.1382, GG ε = 0.4891, p= 0.8064. e) Response probability of Po and VM cells for 1 Hz optical stimulation of L5 cells spanning three orders of magnitude. Probabilities are normalized to maximum responses, laser intensity values transformed to log_10_. Slopes for Po and VM are significantly different (simple linear regression: p= 0.0175). Responses for increasing stimulation intensities were tested by mixed effect analysis on raw data (Mixed-effect analysis with GG correction, followed by Šídák’s multiple comparisons test: intensity: F (3,60) = 10.58, GG ε = 0.4877, p= <0.0001, nucleus: F (1,20) = 3.775, p= 0.0662, interaction: F (3,60) = 1.075, GG ε = 0.4877, p= 0.3729).

**Extended Data Fig. 5.**
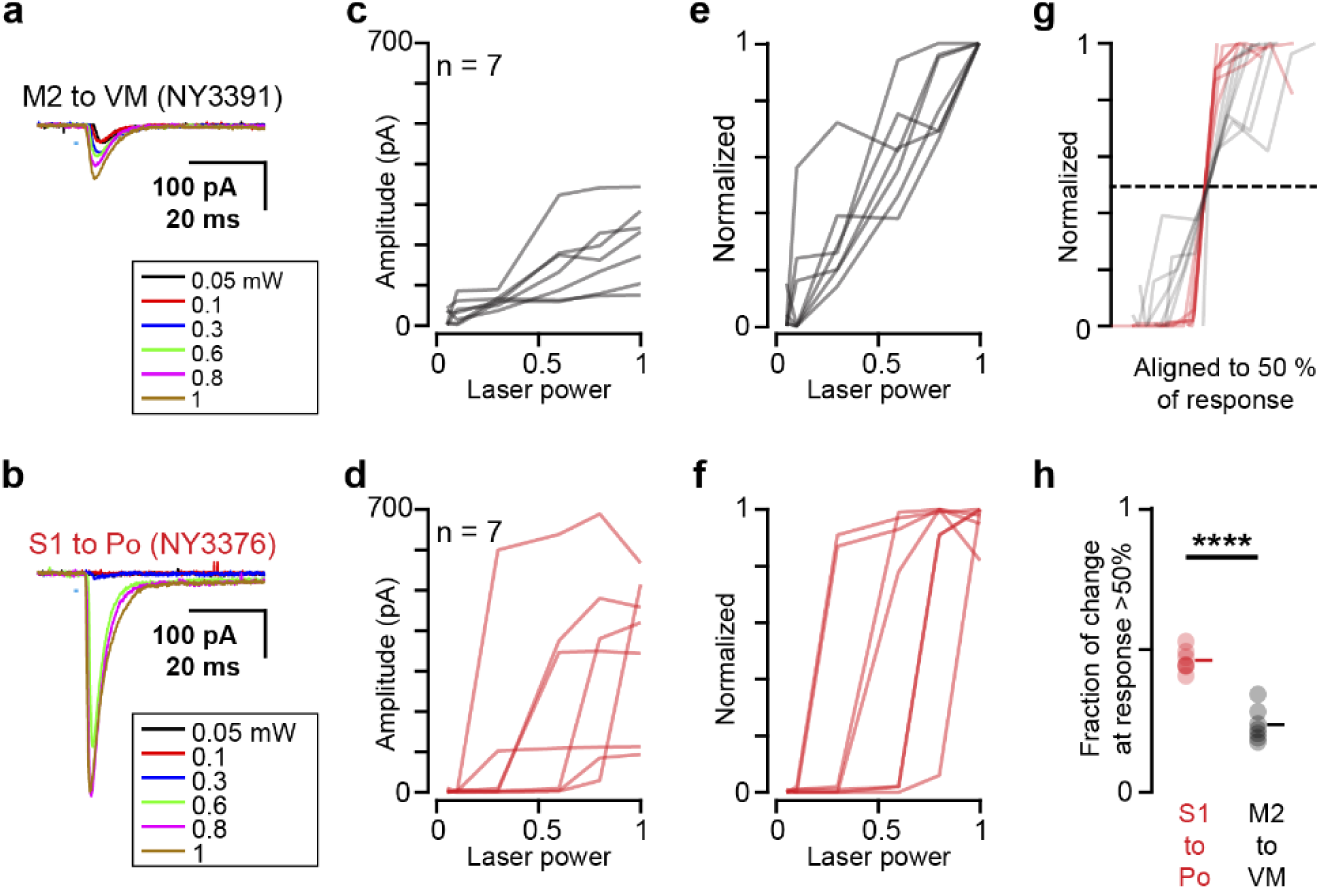
Properties of optogenetically evoked postsynaptic responses *in vitro*. Related to Figure 5. a,b) Example traces of photo-evoked response recorded from a VM (a) or Po(b) neuron upon stimulation of M2/L5 or S1/L5 axons, respectively. Different colors indicate different stimulus intensities. c,d) Group data from the power-series experiment in (a,b) (n = 7). e,f) Data in (c,d) are normalized (See Methods). g) Normalized response of M2/L5-VM and S1/L5-Po pathway in (e,f) were aligned at 50% of max. response and overlaid. h) Statistical comparison of the difference between two responses exceeding 50% of a cell’s maximum response, normalized by the cell’s overall response range (Methods, unpaired t-test, p = 2.11e-6).

**Extended Data Fig. 6.**
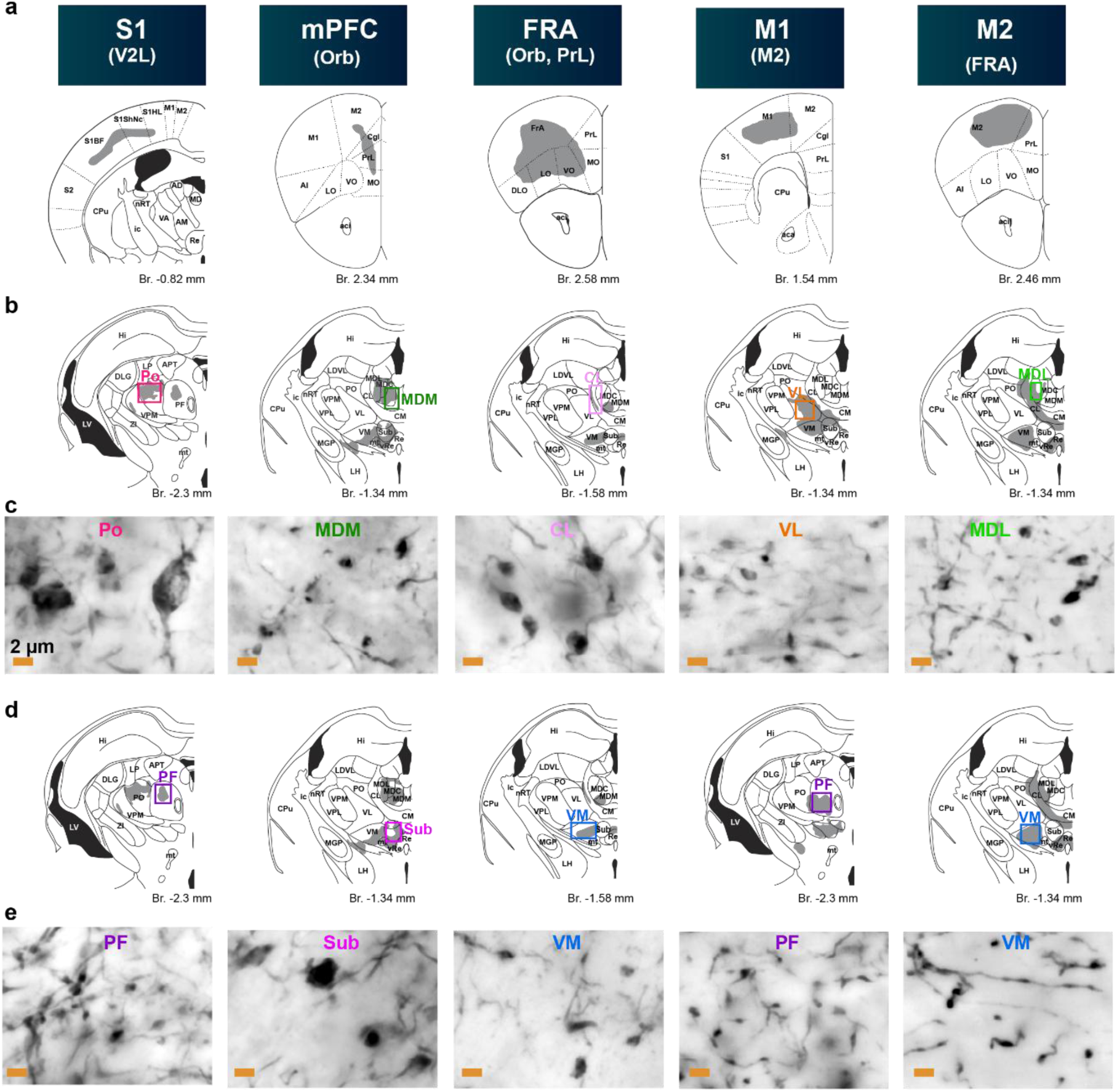
Distribution of L5 terminals with different sizes in the thalamus. Related to Figure 6. a) Schematic drawing of the cortical injection sites in Rbp4-Cre animals. b-e) Thalamic termination fields (b,d) depicting the location (colored boxes) of the high-power light microscopic images (c-e) and nuclear names. Each column represents one experiment.

**Extended Data Fig. 7.**
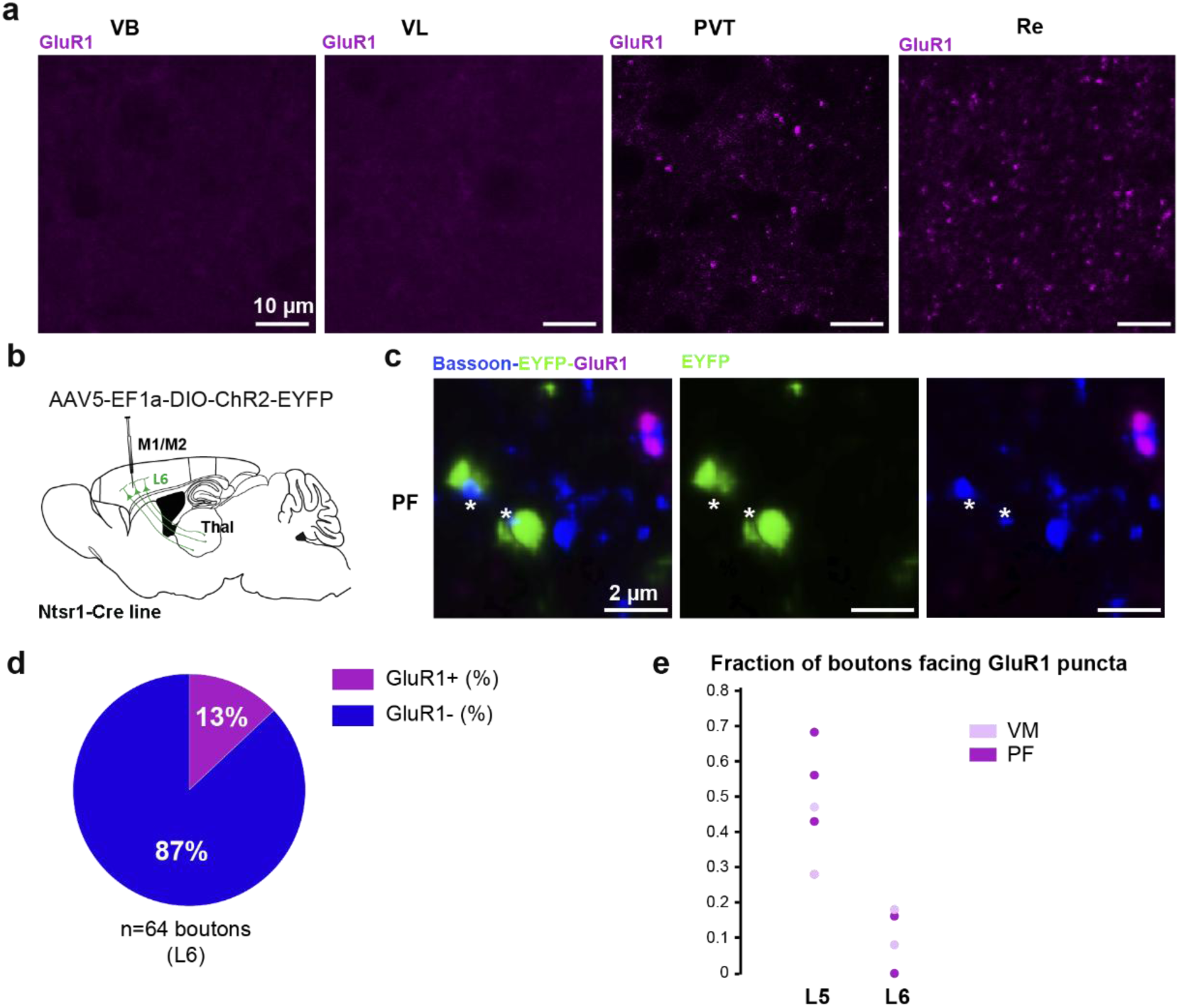
Nuclear distribution of GluR1 receptors and their association with L6 terminals. Related to Figure 6. a) Examples for GluR1 immunostaining in thalamic nuclei receiving no (VB, VL) or small (PVT, Re) L5 terminals. High resolution confocal images. b) Experimental design. c) Examples for GluR1 (magenta)-negative synapses (Bassoon, blue) formed by L6 boutons (EYFP, green) (white asterisks). d) Percentage of GluR1+ and GluR1- L6 synapses in VM and PF (n=64 boutons in n=4 ROI-s in n=2 mice). e) Proportion of GluR1+ L5- and L6-synapses in singe ROIs (L5: 0.45±0.06, n=6 ROI-s in n=3 mice; L6: 0.11±0.04, n=4 ROI-s in n=2 mice).

**Extended Data Fig. 8.**
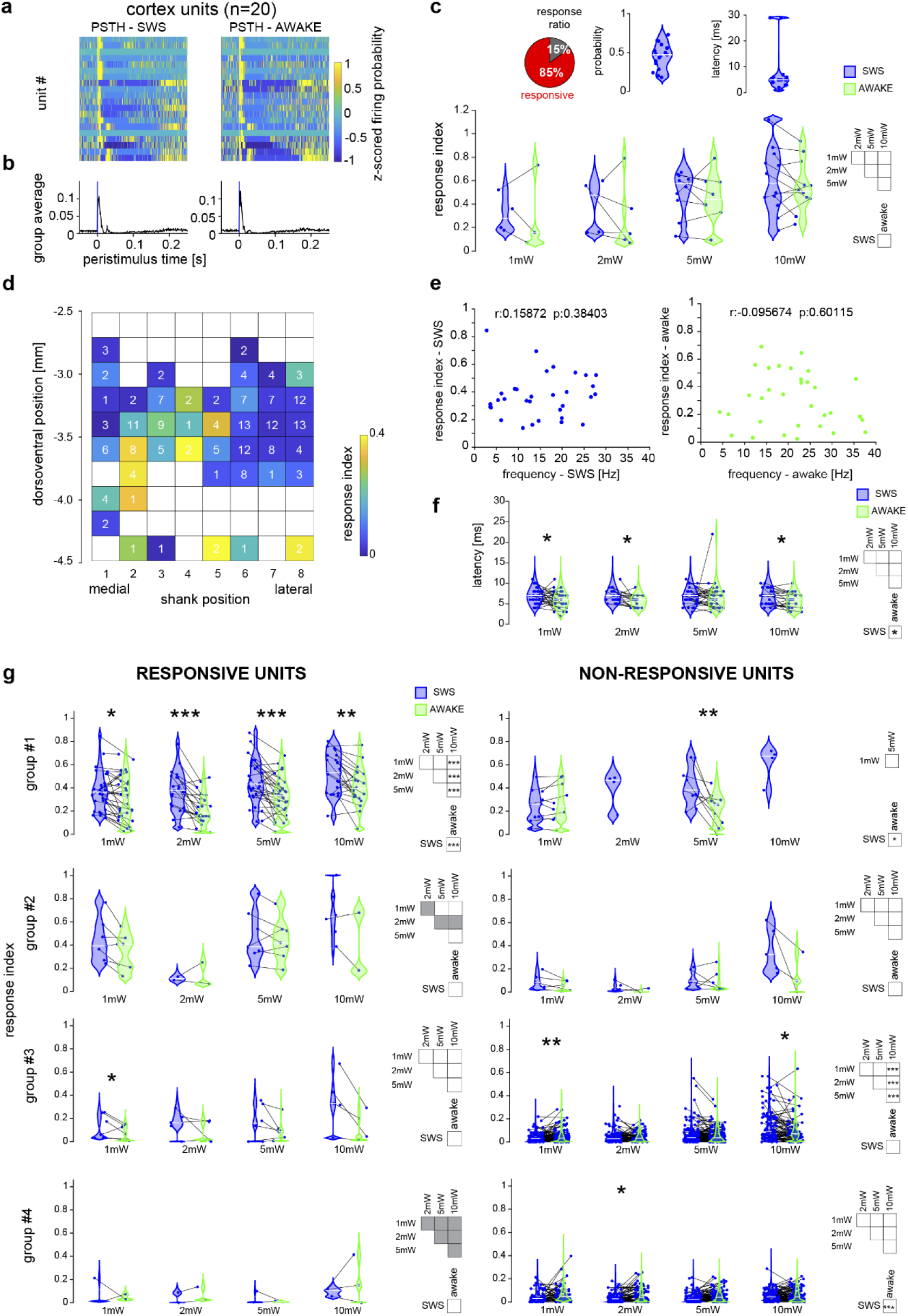
Properties of optogenetically evoked L5 responses in the thalamus of freely behaving animals. Related to Figure 7. a-b) PSTHs of cortical units in both SWS and awake states. c) (Top) Response ratio, probability and latency of L5 cortical responsive units at the injection site. (Bottom) Response indices of cortical responsive units by illumination power. Two-way ANOVA test and post-hoc comparison by Tukey’s HSD test revealed no significant differences. Note that stimulations with all (1 – 10 mW) intensities were carried out only in a subset of cortical units. d) Average response ratio by recording position. Same positional data were used as in Fig. 7. panels f-i. Groups #1-4 are pooled here. Numbers within squares: number of recorded cells at the given location. Note the high variability of response indices at shank 2-4, depth 3.0-3.6 mm even though the densities of L5 terminals were universally high here. e) Spearman correlation between the baseline firing frequency and the response indices for responsive group#1 units, in both SWS (left, blue) and awake (right, green) states. f) Response latency of responsive group #1 units by increasing illumination power, in both SWS and awake states. Two-way RM-ANOVA (intensity: p = 0.039101, state, p = 0.021079, interaction: p = 0.82316) and post-hoc multiple comparison (Tukey’s HSD test), results indicated by asterisks above plots (state effect at each illumination intensity) and in boxes at right (intensity and state effects). g) Response probability indices of responsive (left column) and non-responsive units (right column) for groups #1-4 in both SWS and awake states. Two-way RM ANOVA and post-hoc multiple comparison (Tukey’s HSD test) results indicated by asterisks above plots (state effect at each illumination intensity) and in boxes at right (intensity and state effects), (*: p < 0.05, **: p < 0.01, ***: p < 0.001). Note that the data for responsive group #1 units are identical to Fig. 7 panel l; however, the panels differ in the manner of data organization. Gray boxes indicate very low n (n ≤ 3) in the given group, statistical comparisons for these data are not reported due to insufficient sample size.

Legend for Extended data Video 1.

30 s long video clips of the horizontal wheel running session on day 7 of a representative mouse from the control and the ArchT group (see Fig. 8c, red rectangles) are shown.

## Online Methods

All procedures were conducted in accordance with approvals from the Animal Care and Use Committee of the Institute of Experimental Medicine, HUN-REN, and the Committee for Scientific Ethics of Animal Research of the National Food Chain Safety Office. Experiments were approved by the National Animal Research Authorities of Hungary (PE/EA/877-7/2020 and PE/EA/01173-2/2025). Experiments complied with the 86/609/EEC/2 and 2010/63/EU Directive of the European Council. Mice were housed in temperature- and humidity-controlled facilities on a 12-hour light–dark cycle, with unrestricted access to food and water.

### Anatomical experiments, tracing and immunocytochemistry

For the anatomical investigation of L5 axonal connections in the thalamus from frontal and somatosensory cortical origin the L5 specific Rbp4–Cre mouse line (stock Tg(Rbp4-cre)KL100Gsat/ Mmucd, ID: MMRRC_031125-UCD) was used. Viral tracing injections (AAV5-EF1a-double floxed-hChR2(H134R)-EYFP-WPRE-HGHpA, Addgene 20298-AAV5, 1,87 × 10^13 vg/ml) were stereotaxically targeted to various frontal and somatosensory cortical area (M2: AP +2, ML +0.5, DV −0.7; in case of double injections to M2, M2 anterior: AP: +2.5, ML +0.7, DV, −0.7; M2 posterior: AP +1.5, ML +0.7, DV −0.7; M1: AP +2, ML +2, DV −0.7; FrA: AP +2.7, ML +1.5, DV −0.7; orbitofrontal cortex: AP +2.2, ML +1.2, DV +2; medial prefrontal cortex (mPFC) anterior: AP +2, ML +0.4, DV −1.2; mPFC posterior: AP −1, ML +0.5, DV +0.7 or AP −2, ML −0.5, DV −0.7.; SUM = 56 mice, male, 60-100 days old) Nanoinjections were made by a Nanoliter 2020 Injector (WPI, Sarasota, Florida, 200 nl at 1 nl/s) After 3 to 6 weeks of survival animals were deeply anaesthetized with ketamine/xylazine (ketamine, 83 mg. kg−1, Produlab Pharma, 07/01/2302; xylazine, 3.3 mg kg−1, Produlab Pharma, 07/03/2303) and perfused through the heart first with saline for 2 min. (∼50 ml) then with 200 ml, 4% paraformaldehyde (PF) (TAAB Laboratory, P001) and 50 µm thick coronal slices were cut with a vibratome (Leica VT1200S). Following extensive washes with phosphate buffer (0.1 M PB, Molar Chemicals, 16050 and 02410) and 30 min. incubation with 3% serum-bovine albumine (BSA, 3%, Sigma-Aldrich, A2153) in PB, also containing 0.3 % Triton-X (0.1-0.5%, Sigma-Aldrich, T8787), the EYFP signal in the axons was amplified by immuno-labeling using chicken anti-GFP antibody (1:5000, Thermo Fisher Scientific, A10262), followed by goat anti-chicken-Alexa 488 antibody (1:500, Thermo Fisher Scientific, A-11039). To highlight the borders of higher-order thalamic nuclei, rabbit anti-calbindin antibody, (1:2,000, Swant, CB-38a) then donkey anti-rabbit Cy5, 1:500, Jackson, AB_2340607) was used. Slices were mounted in Vectashield mounting medium (#H-1000-10).

For the quantification of the anatomical observation (Fig. 6 and Extended Data Fig. 6), L5 axonal EYFP signal was visualized for light microscopical observations. Rabbit anti-GFP antibody (1:10000, ThermoFisher, #A-11122) was used followed by biotinylated goat anti-rabbit (1:300, Vector Laboratories, BA-1000) and avidin biotinylated horseradish peroxidase complex (ABC, 1:300, Vector Laboratories, PK-4000). Nickel-intensified 3,3′-diaminobenzidine (DABNi, bluish-black reaction product, DAB: Sigma-Aldrich, D5637) was used as a chromogen. For quantification 100 terminals with the largest diameter have been selected in each nuclei.

### Quantification of terminal size in confocal images

For confocal microscopical observations Nikon A1R, for light microscopy; Zeiss Axio Imager M1 microscope coupled to an Axiocam HrC digital camera was used. When necessary, brightness and contrast were adjusted using Adobe Photoshop (Adobe Systems, San Jose, CA, USA) applied to whole images only. To quantify the size of layer 5 boutons in the thalamus in the S1/L5-Po and M2/L5-VM pathways (Figure 1), a subset of the Rbp4-cre mice (see above) injected into the primary sensory cortex (n=3 animals) and into the secondary motor cortex (n=2 animals) were used. After three weeks of survival period, mice were perfused, and immunocytochemical reactions were performed as above. Brain sections containing the VM and the Po thalamic nuclei were investigated in the confocal microscope (Nikon C2). Confocal Z-stacks (60x objective oil immersion, NA = 1.4;, image dimensions: 70 x 70 × 10 µm, pixel size: x, y: 87 nm, z: 125 nm) were taken then were deconvolved using Huygens Pro software. Image analysis was done using Fiji ImageJ (NIH, Bethesda, MD, USA). Boutons were first located and marked and then outlined by hand at their biggest cross-section. Statistical analysis was done in GraphPad Prism 10.

### GlUR1 immunostaining

To label L5 or L6 terminals, virus injections were performed in adult male Rbp4-Cre (n = 3 mice) and Ntsr1-Cre (n = 2 mice) mice, respectively. AAV5.EF1a.DIO.hChR2(H134R)-eYFP.WPRE.hGH virus (based on Addgene plasmid #20298, UNC Vector Core) was injected bilaterally into the neocortex (M2; AP: 2 mm, ML: 1 mm, DV: 0.8 mm; 200 nl at 1 nl/s). Mice were perfused with 2% paraformaldehyde in 0.1 M PB buffer. Coronal sections (50 μm thick) were cut using a vibratome. For antigen retrieval, sections were incubated in 2 mg/ml pepsin dissolved in 0.2 M HCl at 37°C for 0.5–5 min, with incubation times determined on control samples. The reaction was stopped in ice-cold 0.1 M PB. Sections were then incubated for 30 min in blocking solution containing 10% normal donkey serum (Jackson ImmunoResearch, #017-000-121). Subsequently, slices were incubated overnight at room temperature in 0.1 M PB containing guinea pig anti-GluR1 antibody (1:200), mouse anti-Bassoon antibody (1:5000), and, for labeling L5 or L6 terminals, chicken anti-EYFP antibody (1:5000, Thermo Fisher, #A10262). Secondary antibodies (donkey anti-guinea pig Alexa Fluor 488, #703-545-155; donkey anti-mouse Cy5, Jackson ImmunoResearch, #AB_2340820; goat anti-chicken Alexa Fluor 488, Thermo Fisher, #A-11039) were applied at 1:500 dilution for 2 h at room temperature. Slices were mounted in Vectashield.

Images were acquired using a Nikon AR1 confocal microscope. To analyze the GluR1 content of different thalamic nuclei, images were acquired with a 20× objective (0.2 µm/pixel). Regions of interest (ROIs; 50 × 50 µm) were selected, and background intensity was normalized across images before quantitative analysis. The number of GluR1 puncta was determined in Fiji using a custom-made script. To analyze the GluR1 content of L5 or L6 synapses, virally labeled EYFP+ structures in VM or PF were first identified as synaptic terminals based on Bassoon immunoreactivity in z-stacks acquired with a 60× objective (0.04 µm/pixel, z-step size: 0.1 µm) following deconvolution with Huygens Pro software (Scientific Volume Imaging, Hilversum, The Netherlands). Close appositions between Bassoon and GluR1 puncta were determined for each terminal using slice view in NIS-Elements. Statistical analysis was performed with GraphPad Prism 10.

### Electron microscopy

M1/M2 cortex (AP: 2 mm; ML 1-2 mm, V: 0.6 mm) of adult male Rbp4-Cre (n=2) or Ntsr1-Cre (n=1) mice were injected with AAV-EF1a-DIO-ChR2-EYFP virus. In addition, in a Thy1-EYFP mouse M1/M2 was injected with Phaseolus Vulgaris Agglutinin (PHAL, n=1). Two weeks post-surgery, mice were perfused with 4% paraformaldehyde and 0.1% glutaraldehyde in PB (0.1 M). Coronal sections (50 μm thick) were cut with vibratome. To permeabilize the membranes sections were incubated in sucrose (30%) overnight, followed by freeze-thawing over liquid nitrogen. For visualizing labeled fibers in case of virus injections, sections were incubated with rabbit anti-GFP antibody (1:2000, ThermoFisher, #A-11122) overnight. For staining, Phal labeled fibers, rabbit anti-phaseolus vulgaris agglutinin antibody (1:10 000; Vector, #AS-2300-1) was used. This step was followed in both cases by incubating the sections with biotinylated b-SP donkey anti-rabbit antibody (1:300; Jackson, #AB_2340593). Sections were incubated with ABC complex (1:300) for 2 h. The immunostaining was developed with DABNi. Sections were treated with OsO4, dehydrated in ethanol and propylene oxide, and embedded in Durcupan (Sigma Aldricht, #44610). During dehydration, sections were treated with 1% uranyl acetate in 70% ethanol. Selected blocks were reembedded, 60-nm-thick ultrathin sections were cut with ultramicrotome. Sections were mounted on copper grids. Electron micrographs were taken with Megaview digital camera running on HITACHI 7100 electron microscope. Reconstruct™ software was utilized for 3D reconstruction. Cell membranes of pre- and postsynaptic structures, postsynaptic densities (PSD) and mitochondria in the boutons were reconstructed. For measurements, Fiji software was used on raw pictures. Minor diameters of postsynaptic dendrites were measured in 3 non-consecutive sections and averaged, spine neck diameter was calculated on 3 sections and averaged. Bouton and spine head volume, PSD area and number of mitochondria were calculated in boutons where sections containing the full extent of the given structure were preserved, with a maximum of two consecutive sections missing in the middle of the structure and the ultrastructure of the tissue was appropriate. Dendritic structures were categorized as small appendages when their minor diameter (measured and averaged on three non-consecutive sections) did not reach 300 nm. Structures were regarded as stubby spines when their minor diameter reached 300 nm and the height/width rate at the base (measured on 3D reconstructed dendrites) exceeded 0.5. Mushroom spines were defined as large headed spines (minor diameter > 300 nm, measured and averaged on three non-consecutive sections), with well-defined neck region. PSD complexity index was calculated as in^63^. The presence or absence of the spine apparatus was determined in spine heads whose ultrastructure was well preserved throughout the entire head volume. In some cases, a single bouton targeted multiple postsynaptic structures, resulting in two values being assigned to the same bouton. For each of these parameters only those boutons were included in the analysis where the criteria, as defined, were fully and clearly visible in the entire series. All statistical analyses were performed using GraphPad Prism 10.

### In vitro 2P imaging and uncaging experiments

#### Slice preparation and in vitro patch-clamp recordings

Male or female calb1/ires2cre mice (n=34, 53-123 days old, median age =72) were deeply anaesthetized with ketamine/xylazine (as above) and injected with AAV5-hSyn-DIO-mCherry (Plasmid #50459, Addgene; titer ≥ 1.4 × 10^12^diluted with sterile saline 1:5) at the following coordinates in mm: Bregma -1.2, lateral 1.0, ventral 4.0 to target the ventromedial nucleus of the thalamus. The calb1/ires2cre line was used, as the calcium-binding protein, calbindin is a known marker for the so-called thalamic matrix nuclei, among others, the VM (REF). Mice were sacrificed 10-79 days later (median post injection days = 33) to prepare coronal slices (300 µm) according to previously established methods^73,74^. Briefly, animals were deeply anaesthetized with 5% isoflurane and quickly perfused through the heart with ice-cold cutting solution containing (in mM): sucrose 220, NaHCO3 28, KCl 2.5, NaH2PO4 1.25, CaCl2 0.5, MgCl2 7, glucose 7, Na-pyruvate 3, and ascorbic acid 1, saturated with 95 % O2 and 5 % CO2. The brain was quickly removed and slices were prepared in cutting solution using a vibratome (Vibratome, St. Louis, MO, or Leica VT1000A, Leica Biosystems GmbH, Nussloch, Germany). Slices were incubated in a submerged holding chamber in artificial cerebrospinal fluid (ACSF) at 36 °C for 30 min and then stored in the same chamber at room temperature. For recording, slices were transferred to a custom-made submerged recording chamber under the microscope where experiments were performed at 32-35 °C in ACSF containing (in mM): NaCl 125, KCl 3, NaHCO_3_ 25, NaH_2_PO_4_ 1.25, CaCl_2_ 1.3, MgCl_2_ 1, glucose 25, Na-pyruvate 3, and ascorbic acid 1, saturated with 95% O_2_ and 5% CO_2_. Cells were visualized using an Olympus BX-61 or a Zeiss Axio Examiner epifluorescent microscope equipped with differential interference contrast optics under infrared illumination and water immersion lens (60X, Olympus or 63X, Zeiss). First, the injected thalamic region with densely labeled mCherry expressing neurons was identified with 2P imaging at 810 nm. Cells were patched in this area under oblique or Dodt contrast illumination to perform current-clamp whole-cell recordings from the somata using a BVC-700A (Dagan, Minneapolis, MN) amplifier in the active ‘bridge’ mode, filtered at 3 kHz and digitized at 50 kHz. Patch pipettes (2-6 MΩ) were filled with a solution containing (in mM): K-gluconate 134, KCl 6, HEPES 10, NaCl 4, Mg_2_ATP 4, Tris_2_GTP 0.3, phosphocreatine 14 (pH=7.25), complemented with the Ca^2+^ sensitive dye Oregon Green BAPTA-1 (OGB-1, 100 µM) and Alexa Fluor 594 (50 μΩ; all fluorescent dyes from Invitrogen-Molecular Probes). Series resistance was <30 MΩ. Voltages were not corrected for liquid junction potential (∼10 mV). Cells were kept at -63 ̵ -65 mV. Some experiments were performed in hippocampal CA1 pyramidal cells in slices from the same animals under identical conditions.

Thalamic cells were discarded from further analysis if a) they did not fire rebound bursts, b) their dendrites were severely cut or beaded c) they were located outside of the targeted thalamic area according to *post hoc* anatomical verification.

#### Two photon imaging and glutamate uncaging

Dual galvanometer based two photon scanning systems (Bruker, former Prairie Technologies, Middleton, WI, USA) were used to image neurons and to uncage glutamate^73,75^. Two ultrafast pulsed laser beams (Chameleon Ultra II; Coherent, Auburn, CA, USA) were used, one for imaging (at 920 nm for OGB-1 and at 810-860 nm for Alexa Fluor 594) and the other to photolyze MNI-caged-L-glutamate at 720 nm (Tocris; 10 mM, applied through a puffer pipette with a ∼20-30- µm-diameter, downward-tilted aperture above the slice using pneumatic ejection system (PDES-02TX (NPI, Tamm, Germany)). Laser beam intensity was independently controlled with electro-optical modulators (Model 350-50, Conoptics, Danbury, CT, USA). Emitted light was collected by multi-alkali or GaAsP photomultipliers (Hamamatsu Photonics K.K, Iwata City, Japan). Imaging laser power was kept at relatively low levels to avoid phototoxicity.

Thalamic spines with a head clearly separated from the parent dendrite were chosen for 2P glutamate uncaging. Stimulation of spines was performed by uncaging glutamate ≤0.5 µm lateral to the head of visually identified spines using 0.2-0.5 ms uncaging duration, and laser power was varied to yield EPSPs between 0.2-2 mV peak amplitude. The stimulation was repeated at the same spot 3 times with 490-590 ms interval within each trial, and 2-5 trials were averaged for each condition. The GU spot was at least 1 µm away from the dendrite or other spines. When stimulating spines with a large head, in some cases multiple uncaging locations around the spine head could be tested to find the most effective position. Linescan imaging through the spine head and the parent dendrite was performed at ∼250-1200 Hz with 8 µs dwell time. In some thalamic (but not CA1) neurons the application of MNI glutamate (without 2P illumination) reversibly increased the input resistance and enhanced baseline voltage fluctuation, presumably due to inhibition of GABA_A_ receptors by the compound. While experiments with too large baseline voltage fluctuation were not included in the dataset (see exclusion criteria below) due to distorting EPSPs, the spine Ca^2+^ responses in these cases were qualitatively and quantitatively similar to the rest of the experiments. Recordings were excluded from the analysis if a) the spine head was not clearly separated from the dendritic shaft, b) the uncaging spot was <1 µm lateral from the dendritic shaft, c) baseline fluctuation was large, or d) the spine exhibited signs of photodamage (long-lasting depolarization, permanently elevated spine Ca^2+^ fluorescence and/or drastic spine size or shape change after 2PGU).

For comparison, the same experiments were performed on spines on proximal apical and basal dendrites of CA1PCs (92 ± 6 µm from the soma, n=9 dendrites in 5 cells), with similar results to that reported previously^75^.

Recorded voltage and Ca^2+^ signals were analyzed offline with custom-written macros in IgorPro (WaveMetrics, Lake Oswego, OR, USA) and Python, using averaged traces of 2-5 trials with reliable responses for each condition. Only the first uncaging stimulus was used for analysis, but results were similar when considering all three stimuli within a trial (Extended Data Fig. 3c). All 2PGU stimulation series evoking reliable EPSPs were analyzed. Some Ca^2+^ recordings exhibited a transient laser light artefact during uncaging (before voltage signals began to rise); the affected data points were replaced by the average of the preceding 5 data points. Ca^2+^ signals were expressed as ΔF/F_0_ = (F(t)-F_0_)/F_0_, where F(t) is fluorescence at a given time point and F_0_ is the mean fluorescence during 50 ms preceding the first uncaging stimulus or the depolarizing I_inj_. PMT dark baseline pixel signal was subtracted. Ca^2+^ signal amplitude was measured as the maximum average of 9 consecutive points within 50-70 ms after uncaging. This measurement may slightly overestimate the amplitude and was therefore corrected by subtracting an offset, calculated as the mean of peak amplitude values measured the same way on the baseline period preceding the first stimulus. Ca^2+^ traces displayed in the figures were smoothed with a moving average of 3 data points. Spine-soma distance measurements were performed on 2P maximum projection z-stacks of the dye-loaded neurons using ImageJ (NIH, Bethesda, MD, USA).

Statistical analyses of these data were performed with the Statistica software (14.0.1.25).

#### Post hoc immunochemical labeling of 2P imaged and patch clamp recorded cells

Brain slices with recorded and biocytin filled cells were transferred from the in vitro recording chamber to 4 % paraformaldehyde solution (in 0.1 M phosphate buffer, PB) for overnight incubation. After washing in PB (3×15 min), slices were incubated in primary antibody (all in PB) for mCherry (rabbit 1:3000, BioVision, 5993-100), and in 14 cells, also for the presynaptic active zone protein, Bassoon (mouse, 1:5000, Abcam, Cambridge, UK) overnight, at room temperature. As a second layer, fluorescent molecule conjugated secondary antibodies; goat-anti-rabbit Cy3 antibody (1:500, Jackson, Ely, UK) and goat-anti-mouse-A488, Molecular Probes) were used respectively. Biocytin labeled cells were visualized by streptavidin-Alexa 488 (1:1000, Molecular Probes) or streptavidin-abberior STAR 635P (for STED imaging, 1:1000, Abberior, Göttingen, Germany). Slices were mounted on slides with vectashield for confocal (Nikon A1R) or with SlowFade™ Diamond (ThermoFisher Sccientific, Waltham, MA USA) for STED microscopical observations with the Abberior Facility Line (abberior GmbH, Göttingen, Germany) STED microscope.

### In vivo juxtacellular recording and labeling

#### Surgery

For juxtacellular experiments adult male mice (n=22, 68-172 days old, median age =102) from the L5 specific Rbp4–Cre mouse line (see above) were used. For viral injection mice were anesthetized with an intraperitoneal injection of ketamine-xylazine (see above) and placed inside a stereotactic apparatus. AAV5.EF1a. DIO.hChR2(H134R)-eYFP.WPRE.hGH (based on Addgene plasmid 20298, UNC Vector Core, same as used for anatomical investigations) virus was injected in the right-side of the neocortex (200 nl, 2 nl/s) using borosilicate glass capillaries (d = 20 μm) . Stereotaxic coordinates were the following (S1: AP;-0.7 mm ML; 3 mm from Bregma, DV; 0.7 mm from the brain surface, M2: AP; 2.2 mm, ML; 0.8 mm, DV; 0.7 mm).

#### Juxtacellular recording and labeling

Depth of anesthesia was monitored throughout the experiment by cortical LFP recording. Cortical LFP was recorded in the M2 or S1 cortex with bipolar tungsten electrode (shanks at the surface and ∼ 1 mm deep from the cortical surface, ∼1 MΩ, FHC, Bowdoin, ME, USA). Signals were filtered (0.1 Hz–5 kHz), amplified (Supertech BioAmp, Supertech, Pécs, Hungary), sampled at 20 kHz (micro 1401 mkii, CED, Cambridge, UK), and recorded by Spike2 7.0 software (CED, UK). The local field potential (LFP) recorded in the M2 or S1 cortices displayed slow oscillations (1–2 Hz) that are characteristic of the stages III-3-4 described by^76^. If this cortical slow oscillation showed signs of desynchronization together with whisker twitches (approx. after 40 min- 1 hour from last anesthetic injection) an additional dose of anesthetic (ketamine, 28 mg kg−1; xylazine, 1.1 mg kg−1) was applied intramuscularly.

Single unit activity in the thalamus was recorded by glass microelectrodes (in vivo impedance of 20–40 MΩ) pulled from borosilicate glass capillaries (1.5-mm outer diameter, 0.75-mm or 0.86-mm inner diameter, Sutter Instrument) and filled with 0.5 M K+-acetate and 2% neurobiotin (Vector Laboratories, SP-1120). Electrodes were lowered by a micromanipulator (Scientifica, UK) to the target area (VM: from Bregma: AP −1.3 mm, ML +0.9 mm, from brain surface DV −3.8 to 4.3 mm, Po: from Bregma: AP −1.8 mm, ML +1.5 mm, from brain surface DV −2.8 to 3.5 mm). Neuronal signals were amplified by a DC amplifier (Axoclamp 2B, Axon Instruments/Molecular Devices), further amplified and filtered between 0.16 Hz and 5 kHz by a signal conditioner (LinearAmp, Supertech) and recorded by Spike2 7.0. Juxtacellular labeling of the recorded neurons was done as described previously^55^.

#### In vivo optogenetic activation

The skull was thinned above the M2 of the right hemisphere, where the optic fiber (100 μm core, 0.22 NA) was positioned (S1: AP -0.7 mm, ML 3 mm, M2: AP; 2.2 mm, ML; 0.8 mm from Bregma). Laser beam was generated by a 473-nm DPSS laser (Laserglow Technologies, Toronto, Canada). Laser power at the optic fiber tip was measured before and after each experiment with a photometer (Thorlabs) For each unit the baseline activity was recorded for 300 seconds, then five stimulus trains of 10 stimuli (5 ms long) at 1, 4, 10 and 20 Hz, generated by Spike2 7.0 software (CED) were applied.

#### Histology

At the end of experiments, mice were perfused (same as above, see above in Methods, Anatomical experiments section) and coronal, 50 μm thick sections were cut with a vibratome. After pretreatment with BSA and TritonX (see above) the EYFP fluorescent signal in L5 axons was intensified with chicken anti-GFP antibody (1:5,000, Thermo Fisher Scientific, A10262), followed by goat anti-chicken-Alexa 488 antibody (1:500, Thermo Fisher Scientific, A-11039) Neurobiotin content of the juxtacellularly labeled cells was visualized by Cy3-streptavidin (1:500, Jackson, 434315), higher-order thalamic nuclei were labeled with rabbit anti-CB antibody (1:2,000, Swant, CB-38a) followed by donkey anti-rabbit Cy5, 1:500, Jackson, AB_2340607). Sections were mounted on glass slides (mounted with Vectashield) and observed with a Nikon AR1 confocal microscope. When necessary, brightness and contrast were adjusted using Adobe Photoshop CS2 (Adobe Systems, San Jose, CA, USA) applied to whole images only.

#### Data analysis

We have excluded 2 Po and 2 VM units from the Po (n=9) and VM (n=17) database based on the criteria that their response probability at the maximum intensity photoactivation (10 mW, 1 Hz) was still very low < 0.15 or their latency was longer than 20 ms (also 1.5 SD larger that the population average), making these data unreliable for further analysis. All statistical analysis were performed using GraphPad Prism 10. Group data of probabilities and latencies at different stimulation intensities were compared by 2-way repeated measures (RM) ANOVA with Geisser-Greenhouse correction to adjust for violations of sphericity/homoscedasticity. Latency data for different frequencies, instead of repeated measures ANOVA which cannot handle missing values, data were analyzed with a Mixed-effects model (with Geisser-Greenhouse correction). This method is using a compound symmetry covariance matrix, and is fit using Restricted Maximum Likelihood (REML). In the absence of missing values, this method gives the same P values and multiple comparisons tests as repeated measures ANOVA. In the presence of missing values (missing completely at random), the results can be interpreted like repeated measures ANOVA.

### Laser-scanning photostimulation (LSPS) and repetitive or power-series stimulation of thalamic axons ex vivo

#### Slice preparation and in vitro patch-clamp recordings

Rbp4-Cre mice (male) were injected with AAV5.EF1a.DIO.hChR2 (E123T/T159C). EYFP.WPRE.hGH (Addgene35509) in M2 or S1 (AP +2.2 and +1.5, ML 0.8 for both; N = 2 per group). Mice were euthanized 3–5 weeks after virus injection (age range at the time of recording: 8½–9½ weeks), and coronal brain slices (300 μm) were prepared using a vibratome (VT1200S, Leica) in an ice-cold choline-based cutting solution (composition, in mM: 25 NaHCO3, 1.25 NaH2PO4, 2.5 KCl, 0.5 CaCl2, 7 MgCl2, 110 choline chloride, 11.6 sodium L-ascorbate, and 3.1 sodium pyruvate). Slices were then transferred to artificial cerebrospinal fluid (ACSF; composition, in mM: 127 NaCl, 25 D-glucose, 2.5 KCl, 1 MgCl2, 2 CaCl2, and 1.25 NaH2PO4) for 30 min at 34 °C and then for ≥1 h at room temperature (∼21 °C) before recording. Whole-cell recordings were performed using an upright microscope (BX51WI, Olympus) equipped with gradient contrast and epifluorescence optics. Pipettes (∼2.5–4 MΩ) were filled with a cesium-based internal solution containing (in mM): 128 cesium methanesulfonate, 10 HEPES, 10 phosphocreatine, 4 MgCl2, 4 ATP, 0.4 GTP, 3 ascorbate, 1 QX314, and 1 EGTA (pH 7.25, 290–295 mOsm). To visualize neurons after recording, 0.05 mM Alexa Fluor 488 hydrazide and/or biocytin (4 mg/mL) was added to the internal solution. ACSF, oxygenated with 95% O2/5% CO2, was perfused using a pump-driven recirculation system, and the recording temperature was maintained at 32 °C using an in-line temperature controller (TC-324B, Warner Instruments). Data were acquired using Ephus software. Signals were amplified with an Axon Multiclamp 700B (Molecular Devices), Bessel filtered at 4 kHz, and sampled at 10 kHz.

Whole-cell recordings were established from VM or Po thalamic neurons using a 60× objective lens (LUMPlanFI/IR, NA 0.9, Olympus). The objective was then switched to 4× (UPlanSApo, NA 0.16, Olympus). An 8 × 8 stimulation grid (100 um spacing) was placed over the recorded neuron, with the soma centered within the grid. The laser beam (1 ms, 1 mW) was applied pseudo-randomly to each spot in the grid at 1 Hz. The resulting input map was used to determine the location for repetitive or power-series stimulation at a site away from the soma (>250 um from soma)^57^. For repetitive stimulation, laser power was calibrated to the lowest level required to evoke responses at 1 Hz (0.25 mW increments), and stimulation was then applied at four different frequencies (1, 5, 10, and 20 Hz). For power series analysis, the stimulation site was instead activated with different laser powers (0.05, 0.1, 0.3, 0.6, 0.8, and 1.0 mW). A minimum of three sweeps was collected per frequency or intensity. The order of stimulation frequencies or intensities was randomized for each neuron to minimize sequence effects.

For response normalization in Extended Data Fig. 5 we subtracted the minimum response of each cell from its responses at each power, and then divided by that cell’s response range (i.e., the difference between its maximum and minimum responses). The response comparison was performed by calculating the difference between two responses that represented a change exceeding 50% of the cell’s maximum response, and then dividing this value by the cell’s overall response range (i.e., the difference between its maximum and minimum responses). We excluded neurons with response probabilities at maximum laser intensity below 0.2 OR response latencies longer than 1.5 SD of the population response latencies.

### Freely moving silicone probe recording

#### Animals, surgical procedures

Adult female Rbp4-Cre mice (n = 2) were used for chronic, freely moving experiments. For optogenetic manipulation of the M2/L5 pathway, unilateral injections were made into the M2 cortex (coordinates: AP +2, ML +1, DV 0.7 mm) with 150 nl AAV5-EF1a-double floxed-hChR2(H134R)-EYFP-WPRE-HGHpA viral vector (titer: 2.1×10^13 g/nl). For the virus injection, the mice were anesthetized with an intraperitoneal injection of a ketamine-xylazine mixture (4:1) diluted 6-fold in saline, at a dose of 0.01mg/ g body weight.

Chronic silicon probe implantation in the M2 cortex and thalamus was performed five months after virus injection, under isoflurane anesthesia. Mice were positioned in a stereotaxic frame (David Kopf Instruments, Tujunga, US), the scalp was shaved, disinfected with Betadine, and locally anesthetized with lidocaine. Eye protection was provided with Corneregel ointment (Dr. Gerhard Mann Chem-pharm. Fabrik GmbH, Berlin, Germany), and body temperature was maintained using a heating pad. Respiratory rate was continuously monitored, and anesthesia depth (1.5–2% isoflurane) and airflow (1.3–1.5 l/min) were adjusted as needed using a vaporizer (Rothacher Medical GmbH, Heitenried, CH).

The exposed skull was coated with dental adhesive (Optibond XTR, Kerr, Brea, US). Two craniectomies were performed for silicon probe placement: one targeting the M2 cortex (AP +2.2, ML +1, probe model: A1×32-Poly3-10mm-25s-177) and one targeting the VM thalamus (AP -1.3, ML +1, probe model: Buzsaki64L; both probes from NeuroNexus Technologies, Inc., Ann Arbor, MI US). The M2 probe was equipped with an optic fiber (105 μm core, 0.22 NA), positioned 500 μm above the most dorsal recording site, and implanted at DV 1.1 mm (probe tip position). It remained stationary throughout the experiment. The VM probe was initially placed at DV 2.2 mm and advanced daily by 70–280 μm after each recording session. Both probes were mounted on custom adjustable micro-drives and coated with DiI for later histological verification. Craniectomies were sealed with artificial dura (Cambridge NeuroTech Ltd, Cambridge, UK). Ground and reference wires were inserted beneath the occipital bone. Implants were shielded with copper mesh and dental acrylic (Paladur, Kulzer, Hanau, Germany). Before completing surgery, buprenorphine was administered subcutaneously at a dose of 0.045 μg/g body weight.

#### Chronic recordings

Recordings commenced after a 14–21 day recovery and habituation period. Each session began with home cage control recording, followed by optogenetic stimulation epochs: the M2 cortex was illuminated with 3 ms pulses of 473 nm blue light (1–10 mW intensity at the fiber tip) at 1–20 Hz frequency. After home cage sessions, recordings continued in a cheeseboard maze (control and stimulation epochs). Neural activity was captured using an Intan RHD2000 evaluation board (Intan Technologies, Los Angeles, CA US) and digitized at 20 kHz. Animal movement was tracked with a marker-based 3D motion capture system (Optitrack, 4 Flex 13 cameras, NaturalPoint Inc, Corvallis, OR US), with markers on the headstage and implant shield. Twisting of wires and fiber was prevented using an Imetronic movement compensator (Delta Technologies Intl, Marcheprime, France) and a Doric rotary joint (Doric Lenses, Québec, Canada).

#### Histological verification

At the conclusion of experiments, mice were anesthetized with ketamine-xylazine following isoflurane induction and transcardially perfused with saline (2 min) and 4% paraformaldehyde (20 min). Silicon probes were withdrawn prior to perfusion. Brains were extracted, sectioned coronally (50 μm thick slices), and probe tracks were visualized via DiI fluorescence. Viral transfection of M2/L5 neurons was confirmed by imaging antibody enhanced EYFP fluorescence.

#### Data analysis

For each animal, 6–8 recording sessions were analyzed. In the thalamus, sessions spanned dorsoventral positions of 35–840 μm as the probe was advanced after each session. Spike trains were detected, clustered, and sorted using SpyKING CIRCUS (v1.1.0) with a median filter for movement artifact removal. Clusters were manually curated in phy2.0 based on spike waveform averages, autocorrelograms, amplitude distributions, and principal component analysis. Further analyses were performed in MATLAB with custom scripts. Peri-stimulus time histograms (PSTHs) were computed to assess unit responses to cortical optogenetic stimulation. PSTHs were generated for the lowest intensity tested (1 mW at 1 Hz in the home cage; in three sessions, higher intensities – 2, 5, or 10 mW – were used due to lack of response). Spike counts were binned at 1 ms in a ±1 s window around stimuli. Stimuli were delivered in trains of 10–378 pulses, with 1–7 trains per session depending on stimulation parameters. All trains with identical parameters were pooled, but home cage and maze data were analyzed separately. PSTH spike counts were normalized by stimulus number to yield firing probability. Heuristic K-means clustering of z-scored firing probabilities was used for classification of response.

For response metrics (probability, latency), only the first spikes of bursts (defined as ≥10 ms interspike interval, ≤0.25 s duration) and non-burst spikes were analyzed, smoothed with a 10 ms moving average. The response ratio was defined as the fraction of units with response exceeding a surrogate-derived firing probability threshold for more than 3 ms within the first 30 ms after stimulus. Surrogate stimulus trains (n = 500) preserved original inter-stimulus intervals but were randomly placed within the train-preceding control period. The threshold for response detection was set at the 95th percentile of firing probabilities around surrogate stimuli. For cheeseboard maze recordings with insufficient control periods, thresholds from the longest cheeseboard recording of the given session were used.

Response probability index was calculated as the cumulative firing probability above baseline in bins exceeding baseline within the first 30 ms after stimulus. Baseline was defined as the mean firing probability in the last 30 ms before stimulus. Response latency was the time of the first bin with maximal firing probability in the first 30 ms.

Baseline firing frequency was determined prior to the first stimulus train. The dorsoventral position of each unit was calculated from the probe’s depth and the position of the recording site with maximal waveform amplitude.

For statistical analysis, paired comparisons used the sign rank test, while

comparisons across stimulation parameters or recording states used two-way ANOVA with Tukey’s HSD for multiple comparisons.

### Behavioral experiments

#### Surgery

For behavioral experiments we injected 4x 150 nl of AAV5-CAG-FLEX-ArchT-tdTomato (6×10^12 vg/ml; UNC Vector Core; 28305) or AAV5-CAG-FLEX-tdTomato (7,8×10^12 vg/ml; UNC Vector Core; 20306) bilaterally into the M2 (AP: 1 and 2 mm, ML: 0.5, DV: 0.7) in Rbp4-cre male mice and wild type male litter mates of Rbp4-cre mice. 105 mm optic fibers (Thorlabs, FG105UCA; Precision Fiber Optics, MM-FER2007CF-1260; molded with Loctite Eccobond F112 BIPAX) were placed bilaterally above the VM (AP: -1.4 mm, ML: 1.9 mm, DV: 3.8 mm, in a 14° angle) and fixed with two types of dental cement (Sun Medical Super-Bond Universal Starter Kit; Kulzer Paladur). Before behavioral experiments we allowed a 3 weeks for each animal for viral expression and at least 1 week recovery from the second surgery. All animals were handled before behavioral experiments. Animals were between 65 and 145 days old during the experiments.

#### Open field test

For the open field experiments we placed the animals in a 40×36 cm white plastic arena. During the experiment the animals received laser inhibition for a total of 3 minutes (593 nm, 17 mW, 5 sec ON/10 sec OFF). Control and analysis of the experiments were done by Ethovision XT 15 software (Noldus, Netherlands).

#### Place preference test

For the place preference test we put the animals in 20 x 40 cm two chamber box. The two chambers were marked by different patterns on the walls. Between the two chambers the animals could move freely. Experiments were performed on 3 consecutive days, at the same time after dark period started and lasted for 15 minutes each day. Animals received photoinhibition (593 nm, 17 mW, 5 sec ON/10 sec OFF) on the second day in one chamber in a closed loop system. Control and analysis of the experiments were done by Ethovision XT 15 software.

#### Horizontal wheel running training

For the wheel running experiments we used a sound proofed experimental box, equipped with an electronic rotary joint (Imetronic) and a 1×2 fiber optic rotary joint with intensity divider (Doric). Animals had a 3 day period to accommodate to the rotary joint system (see above). During the wheel running experiments animals were put in the experimental box for 1 hour each day, having free access to the horizontal wheel and received no reward for performing the task. During the experiments the animals received photoinhibiton (593 nm, 17 mW, 5 sec ON/10 sec OFF) in a closed loop system, when entering the wheel’s area. Experiments lasted a total of 7 days with no change in the experimental procedure. Real time closed-loop control of the experiments was commanded with Bonsai costume script, analysis were done by Ethovision XT 15 and custom written MATLAB scripts.

#### Anatomy

24 h before perfusion we performed laser induced microglial activation (475 nm, 17 mW, 5 sec ON/ 10 sec OFF, 10 min total duration), tested by Iba1 staining for microglia, helping the localization of optic fiber tips and anatomical proof of sufficient laser output. Mice were perfused with 4% paraformaldehyde (TAAB Laboratory, P001). Coronal sections (50-μm-thick) were cut with a vibratome (Leica VT 1200S). Slices were permeabilized with 0.3% Triton-X and blocked with 3% Bovine Serum Albumin. Calbindin and Iba1 were visualized with a guinea pig anti-Calbindin D28k antibody (1:2,000, Synaptic Systems, 214 004) and rabbit anti-Iba1 antibody (1:2,000, FUJIFILM Wako Pure Chemical Corporation, 019-19741) followed by DyLight 405 Donkey-anti-guinea pig (1:500, Jackson ImmunoResearch, 706-475-148) and Alexa 488 goat-anti-rabbit (1:500, Molecular Probes, A-11008). All reagents were dissolved in 0.1M PB (Molar Chemicals, 16050 and 02410).

Exclusion criteria: 1 animal was rejected for insufficient microglial reaction (possibly because of fractured optic fibers) and 1 animal because of inappropriate optic fiber placement (left and right optic fibers 950 and 1050 µm dorsally from L5 termination fields in the VM).

#### Statistical analysis

Statistical analyses were performed using GraphPad Prism 10.2.3. For Open Field data we used unpaired t-test. To analyze Place aversion data we used two-way ANOVA. Horizontal wheel learning data were analyzed by one between-subject and one within subject variable design (mixed-ANOVA) of the 2-way repeated measures ANOVA panel in Prism 10, followed by uncorrected Fisher’s LSD.

## LIST OF REFERENCES

1. Colgan, L. A. & Yasuda, R. Plasticity of dendritic spines: Subcompartmentalization of signaling. Annu. Rev. Physiol. 76, 365–385 (2014).

2. Sala, C. & Segal, M. Dendritic spines: the locus of structural and functional plasticity. Physiol. Rev. 94, 141–188 (2014).

3. Nakahata, Y. & Yasuda, R. Plasticity of Spine Structure: Local Signaling, Translation and Cytoskeletal Reorganization. Front. Synaptic Neurosci. 10, (2018).

4. Bourne, J. N. & Harris, K. M. Balancing Structure and Function at Hippocampal Dendritic Spines. Annu. Rev. Neurosci. 31, 47 (2008).

5. Harris, K. M. & Kater, S. B. Dendritic spines: Cellular specializations imparting both stability and flexibility to synaptic function. Annu. Rev. Neurosci. 17, 341–371 (1994).

6. Megías, M., Emri, Z., Freund, T. F. & Gulyás, A. I. Total number and distribution of inhibitory and excitatory synapses on hippocampal CA1 pyramidal cells. Neuroscience 102, 527–540 (2001).

7. Holtmaat, A. & Svoboda, K. Experience-dependent structural synaptic plasticity in the mammalian brain. Nature Reviews Neuroscience 2009 10:9 **10**, 647–658 (2009).

8. Nishiyama, J. & Yasuda, R. Biochemical Computation for Spine Structural Plasticity. Neuron 87, 63–75 (2015).

9. Humeau, Y. & Choquet, D. The next generation of approaches to investigate the link between synaptic plasticity and learning. Nat. Neurosci. 22, 1536–1543 (2019).

10. Fan, L. Z. et al. All-optical physiology resolves a synaptic basis for behavioral timescale plasticity. Cell 186, 543 (2023).

11. Ramon y Cajal, S. Histology of the Nervous System. vol. II (Oxford University Press, Oxford, 1995).

12. Guillery, R. W. A study of Golgi preparations from the dorsal lateral geniculate nucleus of the adult cat. Journal of Comparative Neurology 128, 21–49 (1966).

13. Chen, Y. et al. Thalamic activation of the visual cortex at the single-synapse level. Science 391, 1349–1354 (2026).

14. Shepherd, G. M. G. & Yamawaki, N. Untangling the cortico-thalamo-cortical loop: cellular pieces of a knotty circuit puzzle. Nat. Rev. Neurosci. 22, 389–406 (2021).

15. Acsády, L. Organization of Thalamic Inputs. . in The Thalamus (ed. Halassa, M.) 27–44 (Cambridge University Press, Cambridge, 2022). doi:10.1017/9781108674287.003.

16. Rovó, Z., Ulbert, I. & Acsády, L. Drivers of the primate thalamus. Journal of Neuroscience 32, 17894–17908 (2012).

17. Mason, A., Ilinsky, I. A., Beck, S. & Kultas-Ilinsky, K. Reevaluation of synaptic relationships of cerebellar terminals in the ventral lateral nucleus of the rhesus monkey thalamus based on serial section analysis and three-dimensional reconstruction. Exp. Brain Res. 109, 219–239 (1996).

18. Budisantoso, T., Matsui, K., Kamasawa, N., Fukazawa, Y. & Shigemoto, R. Mechanisms underlying signal filtering at a multisynapse contact. J. Neurosci. 32, 2357–76 (2012).

19. Harris, K. M., Jensen, F. E. & Tsao, B. Three-dimensional structure of dendritic spines and synapses in rat hippocampus (CA1) at postnatal day 15 and adult ages: implications for the maturation of synaptic physiology and long-term potentiation [published erratum appears in J Neurosci 1992 Aug;12(8):following table of contents]. Journal of Neuroscience 12, 2685–2705 (1992).

20. Rouiller, E. M. & Welker, E. A comparative analysis of the morphology of corticothalamic projections in mammals. Brain Res Bull 53, 727–41. (2000).

21. Ojima, H. Terminal morphology and distribution of corticothalamic fibers originating from layers 5 and 6 of cat primary auditory cortex. Cerebral Cortex 4, 646–663 (1994).

22. Bourassa, J., Pinault, D. & Deschênes, M. Corticothalamic Projections from the Cortical Barrel Field to the Somatosensory Thalamus in Rats: A Single-fibre Study Using Biocytin as an Anterograde Tracer. European Journal of Neuroscience 7, 19–30 (1995).

23. Feig, S. & Harting, J. K. Corticocortical Communication Via the Thalamus: Ultrastructural Studies of Corticothalamic Projections From Area 17 to the Lateral Posterior Nucleus of the Cat and Inferior Pulvinar Nucleus of the Owl Monkey Indexing terms: cortical layer 5; thalamocortical; modulation. J. Comp. Neurol 395, 281–295 (1998).

24. Guillery, R. W. & Sherman, S. M. Thalamic relay functions and their role in corticocortical communication: generalizations from the visual system. Neuron 33, 163–75. (2002).

25. Kakei, S., Na, J. & Shinoda, Y. Thalamic terminal morphology and distribution of single corticothalamic axons originating from layers 5 and 6 of the cat motor cortex. Journal of Comparative Neurology 437, 170–185 (2001).

26. Hoogland, P. V, Wouterlood, F. G., Welker, E. & Van der Loos, H. Ultrastructure of giant and small thalamic terminals of cortical origin: a study of the projections from the barrel cortex in mice using Phaseolus vulgaris leuco-agglutinin (PHA-L). Exp. Brain Res. 87, 159–72 (1991).

27. Vidnyánszky, Z. et al. Immunocytochemical visualization of the mGluR1a metabotropic glutamate receptor at synapses of corticothalamic terminals originating from area 17 of the rat. European Journal of Neuroscience 8, 1061–1071 (1996).

28. Sampathkumar, V., Miller-Hansen, A., Murray Sherman, S. & Kasthuri, N. Integration of signals from different cortical areas in higher order thalamic neurons. Proc. Natl. Acad. Sci. U. S. A. 118, e2104137118 (2021).

29. Liu, X. -B, Honda, C. N. & Jones, E. G. Distribution of four types of synapse on physiologically identified relay neurons in the ventral posterior thalamic nucleus of the cat. Journal of Comparative Neurology 352, 69–91 (1995).

30. Feig, S. & Harting, J. K. Ultrastuctural studies of the primate lateral geniculate nucleus: Morphology and spatial relationships of axon terminals arising from the retina, visual cortex (area 17), superior colliculus, parabigminal nucleus, and pretectum of Galago crassicaudatus. Journal of Comparative Neurology 343, 17–34 (1994).

31. Jones, E. G. & Powell, T. P. An electron microscopic study of the mode of termination of cortico-thalamic fibres within the sensory relay nuclei of the thalamus. Proc. R. Soc. Lond. B Biol. Sci. 172, 173–185 (1969).

32. Yamawaki, N. & Shepherd, G. M. G. Synaptic Circuit Organization of Motor Corticothalamic Neurons. Journal of Neuroscience 35, 2293–2307 (2015).

33. Negyessy, L., Hamori, J. & Bentivoglio, M. Contralateral cortical projection to the mediodorsal thalamic nucleus: origin and synaptic organization in the rat. Neuroscience 84, 741–753 (1998).

34. Schwartz, M. L., Dekker, J. J. & Goldman-Rakic, P. S. Dual mode of corticothalamic synaptic termination in the mediodorsal nucleus of the rhesus monkey. J. Comp. Neurol. 309, 289–304 (1991).

35. Prasad, J. A., Carroll, B. J. & Sherman, S. M. Layer 5 Corticofugal Projections from Diverse Cortical Areas: Variations on a Pattern of Thalamic and Extrathalamic Targets. Journal of Neuroscience 40, 5785–5796 (2020).

36. Li, Y. et al. Corticothalamic communication for action coordination in a skilled motor behavior. Nat. Neurosci. 29, 660–672 (2026).

37. Lam, N. H. et al. Prefrontal transthalamic uncertainty processing drives flexible switching. Nature 637, (2025).

38. Schmitt, L. I. et al. Thalamic amplification of cortical connectivity sustains attentional control. Nature 545, 219–223 (2017).

39. Guo, Z. V. et al. Maintenance of persistent activity in a frontal thalamocortical loop. Nature 545, 181–186 (2017).

40. Hádinger, N. et al. Region-selective control of the thalamic reticular nucleus via cortical layer 5 pyramidal cells. Nat. Neurosci. 26, 116–130 (2023).

41. Makino, H. et al. Transformation of Cortex-wide Emergent Properties during Motor Learning. Neuron 94, 880–890.e8 (2017).

42. Mimica, B., Dunn, B. A., Tombaz, T., Srikanth Bojja, V. P. T. N. & Whitlock, J. R. Efficient cortical coding of 3D posture in freely behaving rats. Science (1979). 362, 584–589 (2018).

43. Maclachlan, C., Sahlender, D. A., Hayashi, S., Molnár, Z. & Knott, G. Block face scanning electron microscopy of fluorescently labeled axons without using near infra-red branding. Front. Neuroanat. 12, 409601 (2018).

44. Groh, A. et al. Convergence of cortical and sensory driver inputs on single thalamocortical cells. Cerebral Cortex 24, 3167–3179 (2014).

45. Hoerder-Suabedissen, A. et al. Subset of Cortical Layer 6b Neurons Selectively Innervates Higher Order Thalamic Nuclei in Mice. Cereb. Cortex 28, 1882–1897 (2018).

46. Biró, L., Buday, Z., Kóta, K., Lőrincz, S. & Acsády, L. Convergence and Segregation of Excitatory and Inhibitory Afferents in the Paraventricular Thalamic Nucleus. J. Neurosci. 45, e0539252025 (2025).

47. Harris, K. M. & Stevens, J. K. Dendritic spines of CA 1 pyramidal cells in the rat hippocampus: serial electron microscopy with reference to their biophysical characteristics. Journal of Neuroscience 9, 2982–2997 (1989).

48. Bodor, Á. L., Giber, K., Rovó, Z., Ulbert, I. & Acsády, L. Structural correlates of efficient GABAergic transmission in the basal ganglia-thalamus pathway. Journal of Neuroscience 28, (2008).

49. Errington, A. C., Hughes, S. W. & Crunelli, V. Rhythmic dendritic Ca2+ oscillations in thalamocortical neurons during slow non-REM sleep-related activity in vitro. J. Physiol. 590, 3691–3700 (2012).

50. Spreafico, R. et al. Distribution of AMPA selective glutamate receptors in the thalamus of adult rats and during postnatal development. A light and ultrastructural immunocytochemical study. Developmental Brain Research 82, 231–244 (1994).

51. Yuste, R. & Denk, W. Dendritic spines as basic functional units of neuronal integration. Nature 375, 682–684 (1995).

52. Noguchi, J., Matsuzaki, M., Ellis-Davies, G. C. R. & Kasai, H. Spine-neck geometry determines NMDA receptor-dependent Ca2+ signaling in dendrites. Neuron 46, 609–622 (2005).

53. Sabatini, B. L., Oertner, T. G. & Svoboda, K. The life cycle of Ca2+ ions in dendritic spines. Neuron 33, 439–452 (2002).

54. Sherman, S. M. & Guillery, R. W. Functional organization of thalamocortical relays. J Neurophysiol 76, 1367–95. (1996).

55. Pinault, D. A novel single-cell staining procedure performed in vivo under electrophysiological control: morpho-functional features of juxtacellularly labeled thalamic cells and other central neurons with biocytin or Neurobiotin. J Neurosci Methods 65, 113–36. (1996).

56. Groh, A., de Kock, C. P. J., Wimmer, V. C., Sakmann, B. & Kuner, T. Driver or coincidence detector: modal switch of a corticothalamic giant synapse controlled by spontaneous activity and short-term depression. J. Neurosci. 28, 9652–63 (2008).

57. Jackman, S. L., Beneduce, B. M., Drew, I. R. & Regehr, W. G. Achieving high-frequency optical control of synaptic transmission. J. Neurosci. 34, 7704–7714 (2014).

58. Diering, G. H. & Huganir, R. L. The AMPA Receptor Code of Synaptic Plasticity. Neuron 100, 314–329 (2018).

59. Shi, S. H., Hayashi, Y., Esteban, J. A. & Malinow, R. Subunit-Specific Rules Governing AMPA Receptor Trafficking to Synapses in Hippocampal Pyramidal Neurons. Cell 105, 331–343 (2001).

60. Sherman, S. M. & Guillery, R. W. The Afferent Axons to the Thalamus: Their Structure and Connections. in Exploring the Thalamus and its Role in Cortical Functions 77–137 (MIT Press, Cambridge, Massachusetts, 2005).

61. Kopec, C. D., Real, E., Kessels, H. W. & Malinow, R. GluR1 Links Structural and Functional Plasticity at Excitatory Synapses. The Journal of Neuroscience 27, 13706 (2007).

62. Hering, H. & Sheng, M. Dendritic spines: structure, dynamics and regulation. Nat. Rev. Neurosci. 2, 880–888 (2001).

63. Sun, Y., Smirnov, M., Kamasawa, N. & Yasuda, R. Rapid Ultrastructural Changes in the PSD and Surrounding Membrane after Induction of Structural LTP in Single Dendritic Spines. The Journal of Neuroscience 41, 7003 (2021).

64. Lee, H. K. et al. Phosphorylation of the AMPA receptor GluR1 subunit is required for synaptic plasticity and retention of spatial memory. Cell 112, 631–643 (2003).

65. Clascá, F. Thalamic Output Pathways. in The Thalamus (ed. Halassa, M.) 45–70 (Cambridge University Press, Cambridge, 2022). doi:10.1017/9781108674287.004.

66. Shepherd, G. M. G. & Harris, K. M. Three-dimensional structure and composition of CA3-->CA1 axons in rat hippocampal slices: implications for presynaptic connectivity and compartmentalization. J. Neurosci. 18, 8300–8310 (1998).

67. Vos, M., Lauwers, E. & Verstreken, P. Synaptic mitochondria in synaptic transmission and organization of vesicle pools in health and disease. Front. Synaptic Neurosci. 2, (2010).

68. Cserép, C., Pósfai, B., Schwarcz, A. D. & Dénes, Á. Mitochondrial Ultrastructure Is Coupled to Synaptic Performance at Axonal Release Sites. eNeuro 5, ENEURO.0390-17.2018 (2018).

69. Cathala, L., Holderith, N. B., Nusser, Z., DiGregorio, D. A. & Cull-Candy, S. G. Changes in synaptic structure underlie the developmental speeding of AMPA receptor-mediated EPSCs. Nat. Neurosci. 8, 1310–1318 (2005).

70. Sauerbrei, B. A. et al. Cortical pattern generation during dexterous movement is input-driven. Nature 577, 386–391 (2020).

71. Cohen, J. et al. Cortical up and activated states: Implications for sensory information processing. Neuroscientist 15, 625–634 (2009).

72. Sachdev, R. N. S., Ebner, F. F. & Wilson, C. J. Effect of subthreshold up and down states on the whisker-evoked response in somatosensory cortex. J. Neurophysiol. 92, 3511–3521 (2004).

73. Magó, Á., Weber, J. P., Ujfalussy, B. B. & Makara, J. K. Synaptic plasticity depends on the fine-scale input pattern in thin dendrites of CA1 pyramidal neurons. Journal of Neuroscience 40, 2593–2605 (2020).

74. Kis, N. et al. Cholinergic regulation of dendritic Ca2+ spikes controls firing mode of hippocampal CA3 pyramidal neurons. Proc. Natl. Acad. Sci. U. S. A. 121, (2024).

75. Weber, J. P. et al. Location-dependent synaptic plasticity rules by dendritic spine cooperativity. Nat. Commun. 7, (2016).

76. Friedberg, M. H., Lee, S. M. & Ebner, F. F. Modulation of receptive field properties of thalamic somatosensory neurons by the depth of anesthesia. J Neurophysiol 81, 2243–2252 (1999).

